# Voltage-gated calcium channel subunit α_2_δ-3 shapes light responses of mouse retinal ganglion cells mainly in low and moderate light levels

**DOI:** 10.1101/2020.02.10.941807

**Authors:** Hartwig Seitter, Vithiyanjali Sothilingam, Boris Benkner, Marina Garcia Garrido, Alexandra Kling, Antonella Pirone, Mathias Seeliger, Thomas A. Münch

## Abstract

Little is known about the function of the auxiliary α_2_δ subunits of voltage-gated calcium channels in the retina. We investigated the role of α_2_δ-3 (*Cacna2d3*) using a mouse in which α_2_δ-3 was knocked out by LacZ insertion. Behavior experiments indicated a normal optokinetic reflex in α_2_δ-3 knockout animals. Strong expression of α_2_δ-3 could be localized to horizontal cells using the LacZ-reporter, but horizontal cell mosaic and currents carried by horizontal cell voltage-gated calcium channels were unchanged by the α_2_δ-3 knockout. *In vivo* electroretinography revealed unaffected photoreceptor activity and signal transmission to depolarizing bipolar cells. We recorded visual responses of retinal ganglion cells with multi-electrode arrays in scotopic to photopic luminance levels and found subtle changes in α_2_δ-3 knockout retinas. Spontaneous activity in OFF ganglion cells was elevated in all luminance levels. Differential response strength to high- and low-contrast Gaussian white noise was compressed in ON ganglion cells during mesopic ambient luminance and in OFF ganglion cells during scotopic and mesopic ambient luminances. In a subset of ON ganglion cells, we found a sharp increase in baseline spiking after the presentation of drifting gratings in scotopic luminance. This increase happened after gratings of different spatial properties in knockout compared to wild type retinas. In a subset of ON ganglion cells of the α_2_δ-3 knockout, we found altered delays in rebound-like spiking to full-field contrast steps in scotopic luminance. In conclusion, α_2_δ-3 seems to participate in shaping visual responses mostly within brightness regimes when rods or both rods and cones are active.

## Introduction

Voltage-gated calcium channels (VGCC) of the L-, P/Q-, R- and N-Type form multimeric complexes composed of pore-forming α_1_ subunits, modulatory β subunits and auxiliary α_2_δ subunits (Dolphin, 2012). The α_1_ subunits determine the main biophysical and pharmacological characteristics of the VGCC complex (Catterall et al., 2005), β subunits modulate channel properties, while the α_2_δ subunits mainly promote trafficking (Davies et al., 2007) and targeting of the VGCC complexes (Thoreson et al., 2013), e.g. to lipid rafts (Robinson et al., 2011). Thus, α_2_δ subunits increase calcium current densities and participate in establishing calcium nanodomains. Recent studies have shown a role of α_2_δ subunits in synaptogenesis, independent of their association with a calcium channel (Kurshan et al., 2009), as well as in synaptic structuring (Pirone et al., 2014). α_2_δ subunits get cleaved post-translationally into α_2_ and δ parts and re-linked by disulfide bonds: The δ part anchors α_2_δ to the plasma membrane, putatively by a glycosylphosphatidylinositol (GPI) anchor (Davies et al., 2010), whereas the highly glycosylated extracellular α_2_ part links α_2_δ to the α_1_ subunits (Gurnett et al., 1997). Four isoforms of α_2_δ have been described, termed α_2_δ-1 to α_2_δ-4 (Curtis and Catterall, 1984; Leung et al., 1987; Ellis et al., 1988; Klugbauer et al., 1999; Qin et al., 2002).

In vertebrate retina, photoreceptors mainly express VGCC containing the α_1_ subunit Ca_V_1.4 (Morgans, 2001), β_2_ (Ball et al., 2002) and α_2_δ-4 (Wycisk et al., 2006). Loss of α_2_δ-4 leads to an altered b-wave in the electroretinogram and to photoreceptor degeneration (Wycisk et al., 2006), reminiscent to what is observed in β_2_ knockout mice (Ball et al., 2002). The other ribbon synapse-bearing cell types of the retina, the bipolar cells, also expresses α_2_δ-4 in tiger salamander (Thoreson et al., 2013) and mouse (de Sevilla Muller et al., 2013). So far only few studies have investigated the expression of the other three α_2_δ subtypes in the retina. Rat retinal ganglion cells were found to express α_2_δ-1, acting as a receptor for thrombospondins (Eroglu et al., 2009). Expression of α_2_δ-3 has been reported in ON bipolar cells isolated from mouse retina (Nakajima et al., 2009) and evidence from co-immunolabeling suggests widespread expression in photoreceptors, bipolar cells, glycinergic amacrine cells and most cells in ganglion cell layer (Pérez de Sevilla Müller et al., 2015). The physiological significance of α_2_δ subunit function in mouse retina is largely unclear.

In the present study, we used an α_2_δ-3 (*Cacna2d3*) knockout mouse (Neely et al., 2010) to study the functional aspects of α_2_δ-3 in the mammalian retina.

## Materials & Methods

All procedures were carried out at room temperature unless otherwise noted. Chemicals were obtained from Sigma Aldrich unless otherwise noted.

### Animals

For RT-PCR experiments, wild type mice of the C3H strain (Sanyal and Bal, 1973) were used. All other experiments were carried out on B6.129P2-*Cacna2d3^tm1Dgen^*/J mice (The Jackson Laboratory, RRID: IMSR_JAX:005780), where α_2_δ-3 (gene name *Cacna2d3*) has been knocked out by targeted insertion of a LacZ reporter (*Cacna2d3*^+/+^ will be referred to as wild type and *Cacna2d3*^-/-^ as knockout animals, for genotyping protocols refer to The Jackson Lab). Animals of either sex were used for experiments. Mice were kept in a 12/12 hour dark/light cycle and were used for each set of experiments on approximately the same circadian time of day. Animal use was in accordance with German and European regulations and approved by the Regierungspräsidium Tübingen.

### Behavioral tests: Optokinetic drum

The visual performance of wild type and knockout mice was examined on the behavioral level in a virtual arena, based on the optokinetic reflex (OKR). The head movements of the mouse were used as the behavioral read-out. The freely moving mouse was placed on an elevated platform in the middle of the arena consisting of four computer monitors. A projection of a virtual cylinder with rotating black and white stripes was presented on the screens. Using an automated camera-based system (Viewer³, Biobserve) we were able to automatically adjust the spatial frequency of the stripe patterns on each monitor individually, relative to the head position of the animal, so the stripe pattern appeared uniform from the view point of the mouse. The rotating stripe pattern elicited head movements corresponding to the angular velocity of the stimulus, which were detected by the automated camera-based system and analyzed quantitatively by custom-written Mathematica (Wolfram) scripts to determine correct tracking behavior. To examine the complete range of visual acuity, the spatial frequency of the stimulus was varied between 0.014 and 0.5 cycles per degree (cpd), corresponding to bar widths of 36° to 1°. These stimuli were presented at Michelson contrast levels between 0% (equal stripe color) and 98% (black and white stripes) and rotated clock-wise or counter-clock-wise. For our experiments, rotation speed of the virtual drum was kept at 12°/s (Mitchiner et al., 1976; Abdeljalil et al., 2005; Lagali et al., 2008; Benkner et al., 2013).

### RT-PCR (reverse transcription-polymerase chain reaction) of α_2_δ genes on total retinal RNA

Adult (8-9 weeks old) C3H mice were euthanized by CO_2_ and decapitated. Young C3H mice of postnatal days 3, 6, 9, 12 and 15/16 were euthanized by decapitation. Retinas were dissected out in ice-cold RNase-free phosphate-buffered saline (PBS). Remnants of retinal pigment epithelium (RPE) were removed (adult retinas) and both retinas of each animal were put immediately in lysis buffer (RLT, Qiagen or Trizol, Invitrogen). Tissue samples were incubated for several minutes in lysis buffer, followed by strong vortexing. Unlysed tissue pieces were triturated and dispersed by pipetting. RNA was isolated from the lysates with a spin column-based kit, following manufacturer’s instructions (RNeasy Mini Kit, Qiagen). The RNA was eluted in nuclease-free water and RNA concentration and quality was measured on a Nanodrop ND-1000 (Thermo Scientific). All RNA samples had a 260/280 ratio above 1.9, indicating good RNA quality. Afterwards, RNA was stored short-term at −20°C or long-term at −150°C or immediately used in a retrotranscription reaction (ProtoScript® AMV First Strand cDNA Synthesis Kit, New England Biolabs or SuperScript® III First-Strand Synthesis System, Invitrogen). In each reaction, 48-55 ng (adult mouse) or 300-450 ng (developmental time line) of RNA was used as template and retrotranscribed with oligo d(T) primers according to the manufacturer’s instructions. cDNA products were stored at −20°C and used as templates for polymerase chain reaction (PCR) using cDNA-specific primers for all α_2_δ transcripts. Primer details are summarized in Table 1. For negative controls, a reaction with α_2_δ-3 primers was run in parallel with water as template. The PCR reactions contained 0.2 µM forward primer (for), 0.2 µM reverse primer (rev), 200 µM dATP, dTTP, dCTP and dGTP each, 1.5 mM MgCl_2_, 0.125 µl Standard-Taq polymerase (New England Biolabs), 1 µl template cDNA, 2.5 µl 10X reaction buffer (New England Biolabs), nuclease-free water to fill up to 25 µl total volume. The reaction was set up on ice and run in a thermocycler (BioRad), using a touchdown PCR program: 94°C, 3 min; (94°C, 20 sec; annealing temperature, 30 sec; 72°C, 30 sec) x 35; 72°C, 5 min; 4°C, ∞. Annealing temperature for all primer pairs started at 66°C, decreasing by 1°C per cycle until 61°C for the remaining cycles.

**Table 1:**
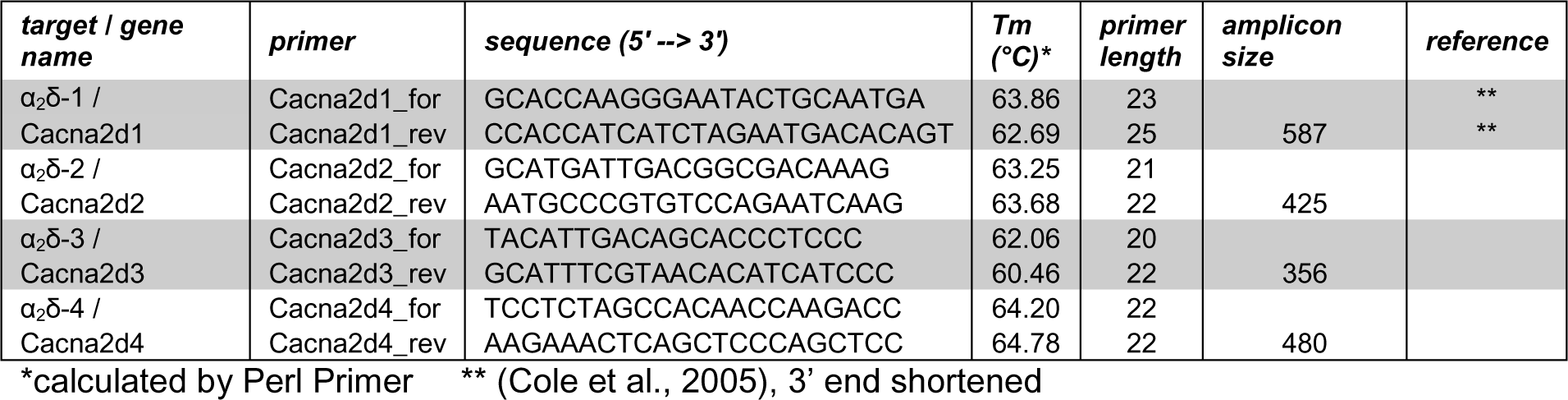
RT-PCR primer details

The PCR products were run on a 1.5% agarose gel, containing 2% midori green (BioZym) or 4% ethidium bromide (Carl Roth) in tris-borate-EDTA buffer (TBE) and visualized using normal UV excitation and filters for ethidium bromide.

*Immunohistochemistry, LacZ reporter expression and horizontal cell mosaic analysis* Adult mice were euthanized by CO_2_ and decapitated. For cryosections, adult (40 days to 7 months) wild type (*Cacna2d3*^+/+^), knockout (*Cacna2d3*^-/-^), and heterozygous (*Cacna2d3*^+/-^) mice were used. For wholemount LacZ stainings, adult (12 to 23 weeks) wild type and knockout animals were used. For horizontal cell mosaic stainings, adult (4-12 weeks old) littermate pairs of wild type and knockout animals were used. Cornea and lens were removed in PBS and eyecups were fixed in 4% paraformaldehyde in PBS for 30 min on ice, or the retina was dissected out, cut to clover-leaf shape, and mounted on a black nitrocellulose filter paper (Sartorius Stedim) before fixation. After washing 3 times in PBS for at least 30 min, retinas were dissected out (wholemount stainings) and retinas/eyecups were cryoprotected by equilibration in a sequence of 10%, 20% and 30% sucrose in PBS.

### Cryosections

The eyecups were transferred to cryo-embedding medium (Tissue-Tek OCT, Sakura Finetek) and frozen on an aluminum block cooled to −150°C. Oriented slices of 12 µm thickness were cut along the dorso-ventral axis in a cryostat (CM3050, Leica), mounted on coated glass slides (Superfrost Plus®, Carl Roth) and dried at 37°C for 30-60 minutes. Slices were stored at −20°C until use. Before proceeding with immunohistochemical staining, slices were thawed and washed 3 times for 10 min in PBS.

Cryoslices were incubated for 12-24 hours at 4°C in a humid chamber with primary antibodies diluted in 10% NGS, 1% BSA, and 0.5% Triton X-100. Antibodies and their dilutions are summarized in Table 2 and their characterization is described below. For negative controls, no primary antibodies were added. Slices were washed 3 times for 10 min in PBS and incubated for 1 hour in a humid chamber with secondary antibodies diluted in PBS, supplemented with 0.5% Triton X-100. Slices were washed 3 times for 10 min in PBS, before being submitted to an X-Gal staining procedure: X-Gal (5% in N,N-dimethylformamide, Carl Roth) was diluted 1:30 in staining solution pre-warmed to 42°C (in mM: 7.2 Na_2_HPO_4_, 2.8 NaH_2_PO_4_, 150 NaCl, 1 MgCl_2_, 3 K_3_(Fe(CN)_6_), 3 K_4_(Fe(CN)_6_); all from Carl Roth, except MgCl_2_, Sigma) and incubated for 12-24 hours at 37°C. Staining was stopped by washing 3 times for 10 min in PBS. Afterwards, slices were washed once in double-distilled water prior to mounting with Vectashield (Vector Laboratories). Images were acquired with an upright epifluorescence microscope (Zeiss Imager Z1), using same settings for stainings and controls, and histograms were adjusted with the min/max function in Zeiss Axiovision software. Images were processed in Adobe Photoshop CS4 and adjusted with the standard auto-contrast function.

**Table 2:** Antibodies and streptavidins used in immunohistochemical stainings on cryosections (c)

| **Antibody name** | **Immunogen** | **Supplier, catalog #, RRID** | **Host species, clonality** | **Dilutions used** |
| --- | --- | --- | --- | --- |
| Calbindin D-28K | calbindin D-28k purified from chicken gut | Swant, 300, AB_10000347 | Mouse, monoclonal IgG1 | 1:500 (c) |
| Calbindin D-28K | recombinant rat calbindin D-28k | Swant, CB-38, AB_10000340 | Rabbit, polyclonal | 1:1000 (w) |
| Neurofilament H | Full length native protein (cow spinal cord) | Abcam, ab4680, AB_304560 | Chicken, polyclonal IgY | 1:1000 (w) |
| Choline Acetyltransferase (ChAT) | Human placental enzyme | Millipore, AB144P, AB_2079751 | Goat, polyclonal | 1:280 (w) |
| Melanopsin | 15 N-terminal extracellular amino acids of mouse melanopsin | Advanced Targeting Systems, AB-N39, AB_1608076 | Rabbit, polyclonal | 1:2800 (w) |
| β-galactosidase | Full length native protein (purified, E. coli) | Abcam, ab9361, AB_307210 | Chicken, polyclonal IgY | 1:500 (w) |
| Anti-mouse/Alexa 488 | IgG (H+L) | Life Technologies, A-21202, AB_10049285 | Donkey, polyclonal IgG | 1:750 (c) |
| Anti-rabbit/Alexa 555 | IgG (H+L) | Life Technologies, A-31572, AB_10562716 | Donkey, polyclonal IgG | 1:400 (w) |
| Anti-chicken/FITC | IgG (H+L) | Jackson ImmunoResearch 703-095-155, AB_2340356 | Donkey, polyclonal IgG | 1:400 (w) |
| Anti-goat/biotin | IgG (H+L) | Life Technologies, D-20698, AB_10375584 | Donkey, polyclonal IgG | 1:400 (w) |
| Streptavidin/Cy5 | nA | Rockland Immunochemicals S000-06, AB_2341143 | nA | 1:200 (w) |

### Wholemount stainings

Retinas were frozen three times at −150°C and thawed at room temperature for better antibody penetration. The retinas were washed three times for 30 minutes in PBS before proceeding with immunohistochemical staining.

Blocking was done in blocking solution (10% normal donkey serum (NDS) or normal goat serum (NGS, for cryoslices), 1% bovine serum albumine (BSA, Carl Roth), 0.5% Triton X-100 in PBS), 0.02% sodium azide for one hour.

One retina of each animal was incubated for 7 days with primary antibodies (see Table 2 and characterization below) diluted in 3% NDS, 1% BSA, 0.5% Triton X-100, 0.02% Na-azide in PBS, gently shaking. The second retina served as a negative control, without addition of primary antibodies. Retinas were washed three times for one hour in PBS after primary antibody incubation. Retinas were incubated with secondary antibodies (Table 2) diluted in 0.5% Triton X-100 in PBS for 24 hours, gently shaking. When biotinylated antibodies were used, retinas were washed three times for 30 min in PBS and incubated with streptavidin-Cy5 in secondary antibody solution for 2-4 hours, gently shaking. Retinas were washed three times for one hour in PBS, cut to clover-leaf shape and flat-mounted on glass slides in Vectashield (Vector Laboratories). Adhesive tape was used as a spacer before adding the coverslip and sealing with nail polish. The stainings and negative controls were evaluated on an upright epifluorescence microscope (Zeiss Imager Z1). The stainings were then visualized on a confocal laser scanning microscope in detail (Zeiss LSM 710). Pinhole sizes were adjusted to the same optical Z-slice thickness for multichannel images.

### Antibody characterization

The mouse monoclonal anti-calbindin antibody (Swant Cat# 300 RRID:AB_10000347) was generated against calbindin D-28k purified from chicken. In western blots, the antibody detects a single ∼28-kDa band in brain extracts of mouse, rat, guinea pig, rabbit and macaque. There is no immunoreactivity of the antibody on hippocampal sections of calbindin D-28k knockout mice (manufacturer’s data sheet, (Celio et al., 1990; Airaksinen et al., 1997)). This antibody is widely used as a marker for retinal horizontal cells in the mouse (Janssen-Bienhold et al., 2009; Phillips et al., 2010) and showed the same labeling pattern as previously described (Haverkamp and Wässle, 2000).

The rabbit polyclonal anti-calbindin antibody (Swant Cat# CB 38 RRID:AB_10000340) was generated against recombinant rat calbindin D-28K. In western blots, the antibody detects a single ∼28 kDa band in brain lysates from mouse, rat, guinea pig, rabbit, macaque and chicken. There is no immunoreactivity of the antibody on cerebellar sections of a calbindin D-28k knockout mice (manufacturer’s data sheet). There is minimal cross-reactivity with calretinin, that gives a ∼30 kDa band in western blots and can also reveal calretinin expression in immunostainings. On the distal inner nuclear layer, the staining however is specific for horizontal cells (Haverkamp et al., 2008; Guo et al., 2009). This antibody is widely used as a marker for retinal horizontal cells in the mouse and showed the same labeling pattern as previously described (Haverkamp and Wässle, 2000).

The chicken polyclonal anti-neurofilament H antibody (Abcam Cat# ab4680, RRID:AB_304560) was generated against purified neurofilaments from bovine spinal cord. Its specificity is confirmed for immunohistochemistry (manufacturer’s data sheet, (Mandadi et al., 2013)). The labeling pattern was the same as in another neurofilament H antibody, selectively labeling axonal arborizations of horizontal cells in the outer plexiform layer (Shelley et al., 2006). Also, in retinal dissociations we observed co-labeled putative horizontal cell axonal arborizations which stained positive for calbindin and the neurofilament antibody.

The goat polyclonal anti-choline acetyltransferase antibody (Millipore Cat# AB144P, RRID:AB_2079751) was generated against human placental enzyme. In western blots, the antibody detects a 68-kDa band e.g. in NIH/3T3 lysate (manufacturer’s data sheet). In the retina, the antibody labels cholinergic ON and OFF starburst amacrine cells and their stratifications in sublaminae 2 and 4 of the inner plexiform layer (Yip et al., 1991; Yonehara et al., 2011).

The rabbit polyclonal anti-melanopsin antibody (Advanced Targeting Systems Cat# AB-N39, RRID:AB_1608076) was generated against the 15 N-terminal extracellular amino acids of mouse melanopsin. Saporin conjugated to this antibody selectively ablates melanopsin-expressing RGC-5 cells and melanopsin-positive ganglion cells in mouse retina (Göz et al., 2008).

The chicken polyclonal anti-β-galactosidase antibody (Abcam Cat# ab9361, RRID:AB_307210) was generated against the full length native protein, purified from E.coli. There is no immunoreactivity in mouse brain samples, that do not transgenetically express β-galactosidase (Scott et al., 2009). We never observed immunoreactivity with this antibody in the retina of wild type animals (which are LacZ negative).

### Horizontal cell mosaic analysis

Horizontal cells and their processes were stained with antibodies against calbindin and neurofilament H in retinal wholemounts. Confocal Z-stacks (5 to 10 Z planes) were acquired with a 10X objective (NA 0.3, 0.83 µm pixel size), covering the retinal wholemount (850 * 850 µm per image), so that all horizontal cells stained were completely visualized including their processes. Overlap between neighboring images was set to 20%. Z-stacks were maximum intensity projected in ZEN 2011 (Zeiss), contrast-adjusted in Fiji (RRID:SciRes_000137, (Schindelin et al., 2012)) and combined with the grid stitch plugin (Preibisch et al., 2009) to yield complete retinal wholemounts. From these, an image part of each of the two dorsal and ventral leaves per wholemount was selected in Photoshop CS4 (Adobe), covering the middle 50% from the optic nerve to outer rim span (see Fig. 5A). The images were analyzed in a custom-written Wolfram Mathematica script to detect cells and compute spacing and mosaic regularity. The neurofilament channel was subtracted from the calbindin channel to produce cleaned-up horizontal cell somata (Peichl and Gonzalez-Soriano, 1994). Images were binarized after adjusting images with dilation/erosion functions. Cell bodies were automatically detected and centroid positions were extracted.

Automatic detection correctly identified between 75 to over 95% of cells, which was then manually corrected for false-positives and false-negatives. With the centroid position information, nearest neighbor distance for each cell could be calculated. Data analysis and statistics were done with Mathematica. For significance testing, the means of the nearest neighbor distances from each individual evaluated image were used (two dorsal and two ventral from each retina).

### Patch-clamp recordings of isolated horizontal cells

#### Retina dissociation procedure

Adult (5-16 weeks old) wild type and knockout mice were anaesthesized by CO_2_, euthanized by cervical dislocation, and retinas were dissected out in CO_2_-independent medium A (per 100 ml): 97 ml HBSS - Ca/Mg free (Biochrom, L2055), 1 ml 10 mM EDTA (final: 0.1 mM, Sigma, E-6511), 1 ml 1 M HEPES (final: 10 mM, Biochrom, L1613), 1 ml 10.000 U/ml Penicillin/Streptomycin (final: 100 U/ml, Biochrom, A2213). Retinas were incubated in a papain solution (medium A, supplemented with 1 mM L-Cysteine (Sigma, C-1276), 20 U/ml papain (Worthington, LS003126)) for 25-40 min at 37°C. Meanwhile medium B was pre-incubated in 5% CO_2_. Medium B (per 100 ml): 10 ml 10X DMEM (Biochrom, F0455), 83.3 ml tissue culture water (PromoCell, C-49998), 2.7 ml 7.5% Sodium bicarbonate (final: 24.11 mM, Biochrom, L1713), 1 ml 1 M HEPES (final: 10 mM, Biochrom, L1613), 2 ml 200 mM L-Glutamine (final: 4 mM, Biochrom, K0283), 1 ml 10.000 U/ml Penicillin/Streptomycin (final: 100 U/ml, Biochrom, A2213). Papain digestion was stopped by replacing the papain solution with DNAse solution (90% Medium B, 10% fetal calf serum (PAA, A15-101), 100 U/ml DNAse I (Sigma, D-5025)) and incubating for 5 min at 37°C. Retinas were centrifuged in a tabletop centrifuge (500 rpm, 3 min) and washed twice with medium B prior to mechanical trituration with fire-polished glass pipettes. In most experiments further gentle dissociation was achieved by shaking the tube for 5-10 min on the slowest setting of a vortexer (Vortex-Genie 2, Scientific Industries). Dissociated cells were seeded on coated coverslips (Ø 12 mm, Gerhard Menzel GmbH, CB00120RA1, coating: 1 mg/ml Concanavalin-A (Sigma, C-7275) in PBS) and kept in medium B at 37°C in 95% O_2_/5% CO_2_ for at least 30 minutes until use. Coverslips were transferred to a recording chamber inside an upright microscope and solution was switched to external solution (see below).

#### Whole-cell patch-clamp recordings

Patch pipettes were pulled from borosilicate glass (Science Products, GB150F-8P), backfilled with internal solution (see below) and had resistances of 4-8 MΩ. Cells were viewed on a digital camera through a 60X water immersion objective (Olympus, NA 1.0) under infrared illumination and horizontal cells were identified by their characteristic morphology (Schubert et al., 2006). Currents were recorded through a patch-clamp amplifier (EPC-10, HEKA) on a computer running PatchMaster (v2×69, RRID:SciRes_000168, HEKA). Internal and external solutions were designed to isolate currents carried by voltage-gated Ca channels.

Internal solution (mM): 110 Cs-methanesulfonate (C1426), 20 TEA (tetraethylammonium)-Cl (T2265), 10 Phosphocreatine-Tris (Santa Cruz, sc-212557), 10 HEPES (4-(2-hydroxyethyl)-1-piperazineethanesulfonic acid, H3375), 5 EGTA (ethylene glycol tetraacetic acid, 03780), 4 ATP (Adenosine triphosphate)-Na (A3377), 0.2 GTP (Guanosine triphosphate)-Na (G8877), 0.51 CaCl_2_ (C5080, free concentration ∼20 nM, calculated with WinMAXC32 v2.50, (Patton et al., 2004)), 4 MgCl_2_ (M2670), pH = 7.2 with CsOH (516988), osmolality 295 mOsm/kg. Chemicals from Sigma, unless noted otherwise.

External solution (mM): 123 NaCl (VWR, 27810.295), 5 KCl (P9333), 10 TEA-Cl (T2265), 10 BaCl_2_ (342920), 1 MgCl_2_ (M2670), 10 HEPES (H3375), 10 D-Glucose (G7528), 1 µM TTX (Biotrend, BN0518), pH = 7.4 with NaOH (S2770), osmolality 305 mOsm/kg. Chemicals from Sigma, unless noted otherwise. For cobalt block experiments, BaCl2 was equimolarly substituted with CoCl2 (Sigma, C8661). Liquid junction potential with these solutions was 14.5 mV (calculated with JPCalc, (Barry, 1994)) and was partially corrected online (set to 10 mV). Post-recording correction was not applied.

Horizontal cell resting membrane potentials were determined by current-clamp to 0 pA directly after establishing whole-cell configuration. Cell capacitances were determined by the Cslow function of the HEKA amplifier. Membrane resistance was generally > 1 GΩ during recording. Cells were held at −60 mV in between voltage protocols. Recording protocols started two minutes after break-in and were interleaved by 30-s pauses in between each protocol. Voltage protocols included (1) maximum current measurements with 50 ms steps to 0 mV, (2) current-voltage (IV) curves with 50 ms steps from −70 to +60 mV and (3) inactivation protocols with 400 ms inactivation steps from −60 to +20 mV and an additional 50 ms test step to 0 mV. Current amplitudes were calculated from the average during the second half (25 ms) of the test steps in each protocol. IV curves were fitted according to the following equation: I = G_max_ (V − V_rev_) / { 1+exp [− (V − V_0.5,act_) / k_act_] }, where I is the current amplitude, G_max_ is the maximum slope conductance, V is the test potential, V_rev_ is the reversal potential, V_0.5,act_ is the half-maximal activation voltage and k_act_ is the slope factor (Ortner et al., 2014). Fitting and parameter calculations were done in SigmaPlot (v12.0, Systat Software).

Recordings were done using p/n leak subtraction (25% magnitude, 4 repetitions, min/max −128 mV/-50 mV). Within each voltage protocol, holding potential was −80 mV. At the beginning of each protocol a 50 ms test pulse to −90 mV was applied. The current difference (non-leak subtracted data) during and after the test pulse was used to calculate input resistance (R_in_) offline using Ohm’s law. Series resistance (R_ser_) was read off the amplifier’s Cslow function after each voltage protocol. Membrane resistance (R_mem_) was calculated by R_mem_ = R_in_ - R_ser_. For maximum current analysis, only voltage protocols were considered that fulfilled two criteria: (1) R_ser_ < 10% R_mem_. (2) mean deviation from zero during 0 mV step > mean deviation from zero during 100 ms before step. Since maximum current amplitude slowly increased with recording time, only maximum current protocols between two and five minutes after break-in were considered for analysis. Only IV curves with clear U-shaped fits were used for analysis. Inactivation data to use for analysis was chosen manually. Data was analyzed with Matlab (Mathworks) and tested for significant differences using Wilcoxon ranksum test.

#### Electroretinography (ERG)

Animals used for ERG recordings were 27-28 days old littermates (n = 5 wild type, n = 4 knockout animals from 5 litters). ERGs were recorded as described previously (Tanimoto et al., 2009). Mice were anaesthetized using ketamine (66.7 mg/kg body weight) and xylazine (11.7 mg/kg body weight). The pupils were dilated and single-flash responses were obtained under scotopic (dark-adapted overnight) and photopic (light-adapted with a background illumination of 30 cd/m^2^ starting 10 minutes before recording). Single white-flash stimuli ranged from - 4 to 1.5 log cd*s/m^2^ under dark-adapted conditions, and from −2 to 1.5 log cd*s/m^2^ under light-adapted conditions. Ten responses were averaged, with interstimulus intervals of 5 s (for −4 to −0.5 log cd*s/m^2^) or 17 s (for 0 to 1.5 log cd*s/m^2^).

#### Multi-electrode array (MEA) recordings: Experimental procedure

Adult (9-12 weeks old) wild type and knockout mice were dark-adapted for at least 4 hours, euthanized by cervical dislocation and retinas were isolated under dim red light in Ringer’s solution containing (in mM): NaCl 110, KCl 2.5, CaCl_2_ 1, MgCl_2_, 1.6, D-glucose 10, NaHCO_3_ 22; bubbled with 5% CO_2_/95% O_2_; pH 7.4. Retinas were mounted on a nitrocellulose filter paper (Millipore) with a 2–3 mm rectangular aperture in the center, transferred to a 60-electrode perforated multi-electrode array (MEA) chamber (60pMEA200/30iR-Ti-gr, Multichannel Systems) and placed ganglion cell-side down on the recording electrodes. The inter-electrode distance was 200 µm, interspersed by holes through which gentle suction was applied. The MEA was put in an upright microscope, where light stimulation was supplied through the microscope condenser by a digital light processing (DLP) projector (PG-F212X-L, Sharp), presenting a diverse set of visual stimuli focused onto the photoreceptors. One batch of stimuli lasted 28 to 40 minutes and was shown at least twice per ambient luminance level. Specific luminance levels were achieved by inserting neutral density filters into the light path. The neutral density (ND) filters (Thorlabs NE10B-A to NE50B-A), had optical densities from 1 (“ND1”, i.e. 10-fold light attenuation) to 5 (“ND5”, 10^5^-fold attenuation). To achieve light attenuation stronger than 5 log units, two ND filters were combined in series. Experiments started at ND8 (‘scotopic’; darkest setting, 10^8^-fold attenuation) and continued over ND6 (‘mesopic’; 10^6^-fold attenuation) to ND4 (‘photopic’; 10^4^-fold light attenuation). A shutter was closed while changing ND filters during the experiment to prevent intermittent exposure to unattenuated light. During the whole experiment the retina was continuously superfused with Ringer’s solution pre-warmed to 38°C by an in-line heating system just before the MEA chamber (33-35°C inside the recording chamber). In this study, total recording times were up to 4.5 hours. Detailed description of the MEA recording procedures can be found in (Reinhard et al., 2014).

### Stimuli

All stimuli were gray-scale images with pixel values between ‘0’ (“black”) and ‘255’ (“white”). The stimulus projector produced an output spanning 3 log units of light intensities (i.e. 1000-fold difference between black (‘0’) and white (‘255’) pixels). We linearized the gamma-function of the projector, so that our background set at ‘128’ corresponded to the middle physical light intensity between ‘0’ and ‘255’. All stimuli were balanced so their mean light intensity over time was ‘128’. Each stimulus was preceded by at least 1 s of uniform background gray (‘128’).

Here, we present data on the following stimuli:

1. Full-field Gaussian white noise (“flicker”) (Chichilnisky, 2001): Each flicker stimulus consisted of five 20-s episodes of high-contrast flicker interleaved with five 20-s episodes of low-contrast flicker. Screen brightness was updated every frame (60 Hz), and drawn from a Gaussian distribution with mean ‘128’ and sigma of 38.4 (high contrast) or 7.68 (low contrast). The flicker stimulus was presented four times per ambient luminance level.
2. Full-field steps: background gray (‘128’) → black (‘0’) → gray (‘128’) → white (‘255’) → gray (‘128’), 2 s per step (±100% Weber contrast).
3. Drifting sine wave gratings: 30 different drifting sinusoidal gratings (combinations of six spatial periods: 100, 200, 500, 1000, 2000, 4000 µm and five temporal frequencies: 0.25, 1, 2, 4, 8 Hz) of full contrast (black: ‘0’ to white: ‘255’).

In addition, moving bars and natural movie stimuli were included in our stimulus batch, but are not discussed here.

#### Light Intensity Measurements

We measured the spectral intensity profile (in µW·cm^−2^·nm^−1^) of our light stimuli with a calibrated spectrophotometer (USB2000+, Ocean Optics). We then transformed the stimulus intensity into equivalents of photoisomerizations per rod per second, assuming dark-adapted rods, as described previously (Munch et al., 2009). Briefly, the spectrum was converted to photons·cm^−2^·s^−1^·nm^−1^, convolved with the normalized spectrum of rod sensitivity (Umino et al., 2008), and multiplied with the effective collection area of rods (0.5 µm²) (Nikonov et al., 2005). The mean light intensity (=background ‘128’) used in this study was 4 Rh*·s^−1^ per rod (ND8, ‘scotopic’), 4·10^2^ Rh*·s^−1^ per rod (ND6, ‘mesopic’) and 4·10^4^ Rh*·s^−1^ per rod (ND4, ‘photopic’).

#### Spike sorting

Data was recorded at 25 kHz with a USB-MEA-system (USB-MEA1060, Multichannel Systems), high-pass filtered (500Hz, 10th-order butterworth filter) and spikes were extracted by thresholding. Spike sorting (assignment of spikes to “units”, presumably individual ganglion cells) was performed semi-manually with an in-house written Matlab (Mathworks) routine, using 2-dimensional projections of e.g. spike amplitude, principal components, Euclidian distance to template waveforms. Quality of each unit was assessed by interspike interval and spike shape variation. Data analysis was based on the spike times of individual units.

#### Spike rate calculation

The instantaneous spike rate of each unit was calculated by convolving the spike train with a Gaussian with sigma = 40 ms and amplitude = .25 √e sigma^−1^ (≈ 10 Hz for sigma = 40 ms) and was used for further analysis in Matlab. Single units will be referred to as individual “cells” in the following text.

#### Determining cell polarity using the spike-triggered average

We calculated spike-triggered averages (Chichilnisky, 2001) in response to the Gaussian flicker by summing the 500 ms stimulus history before each spike during high- or low-contrast episodes. The polarity of the spike-triggered average (STA) was assessed manually for each cell and ambient luminance level; if the first peak was negatively deflected, it was categorized as an OFF cell, if positively deflected it was categorized as an ON cell (Tikidji-Hamburyan et al., 2014).

#### Statistics

Statistical significance was tested with Wilcoxon rank sum tests and data was plotted using Matlab.

## Results

### 1. Optokinetic reflex is present in α_2_δ-3 knockout mice

The optokinetic reflex is a simple reflex behavior in which a moving stimulus is followed by the gaze (“tracking”) of the animal with eye and head movements. In our behavioral setup, the mice were put in a virtual optokinetic drum and presented with stripe patterns of uniform stripe width (Benkner et al., 2013). Stimulus intensity was within the photopic regime. For the analysis in this study, we only considered if animals were able to track the stimulus at any of the tested conditions, to establish whether visually guided behavior is developed in α_2_δ-3 knockout mice. Wild type and α_2_δ-3 knockout animals performed equally well in this simple visually guided behavior (Fig. 1, n = 5 each), with only one knockout animal not showing clear tracking behavior.

**Figure 1:**
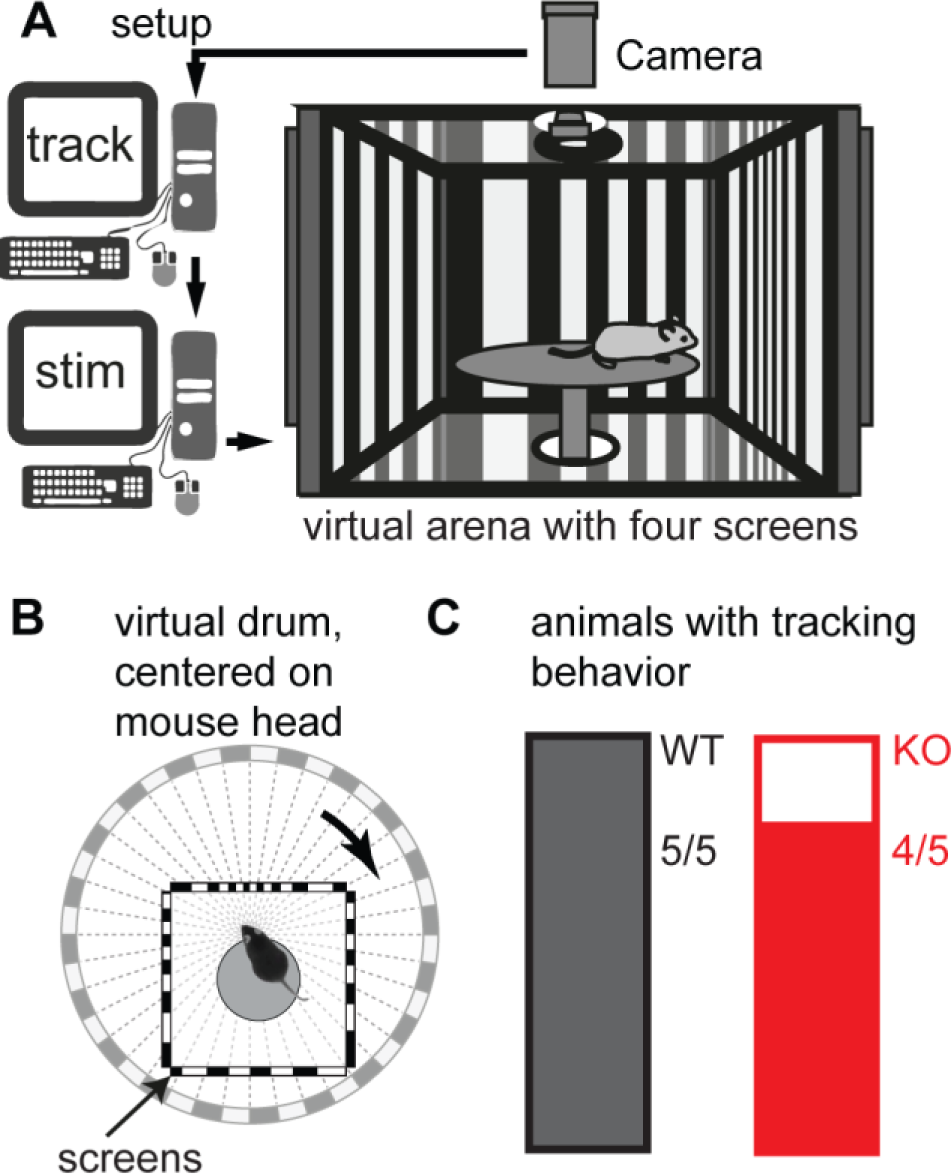
Optokinetic reflex test. (**A**) The “optokinetic drum” virtual arena. Rotating stripe patterns were presented while the mouse was observed from above by a camera. The recorded position of the mouse head was used to adjust stimulus parameters in real time (**B**), circle = virtual drum, square = actual stripe pattern on the screens. (**C**) all 5 wild type and 4 out of 5 knockout animals showed clear tracking behavior, indicating an intact optokinetic reflex in α_2_δ-3 knockout animals.

### 2. All α_2_δ genes are expressed in mouse retina throughout development

We performed RT-PCR to establish which α_2_δ genes are expressed in the retina. Total RNA was isolated from adult C3H mouse retina (n = 3; results were the same with α_2_δ-3 wild type retinas) as well as developing C3H retina (postnatal days 3, 6, 9, 12, 15/16; n = 3 animals each except n = 2 for P6 and P15/16), and RT-PCR was done using cDNA-specific primers for each gene. Negative controls were performed with water as template and never gave a visible band (not shown). All four α_2_δ isoforms were consistently detected at all developmental time points (Fig. 2). The double band for α_2_δ-2 corresponds to different splice isoforms (verified by sequencing).

**Figure 2:**
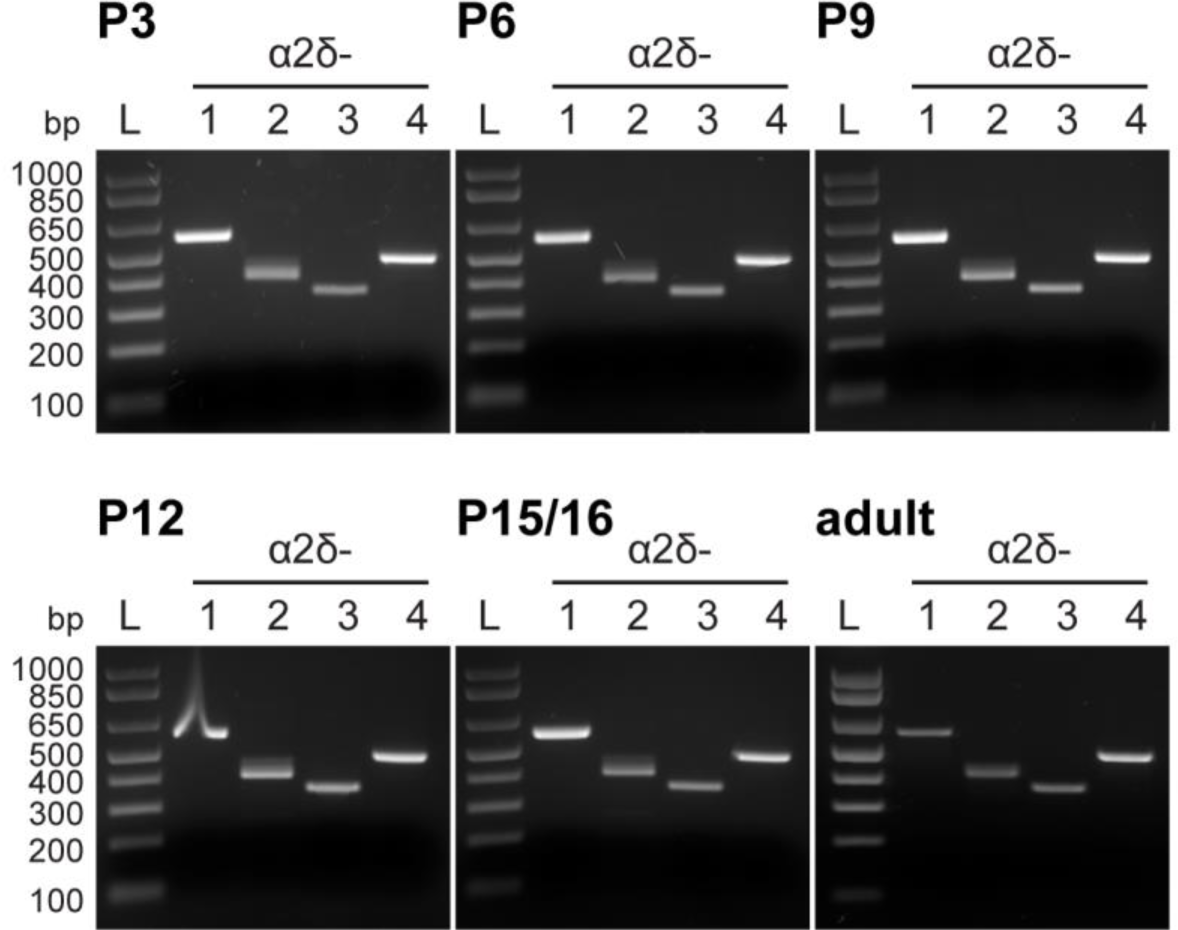
α_2_δ mRNA expression in mouse retina. RT-PCR on total RNA isolated from mouse retina at postnatal day 3 (P3) up to adult. All four α_2_δ transcripts were found throughout the developmental time points tested and in adult mouse retina (n = 3 each, n = 2 for P6 and P15/P16). Amplicon sizes: α_2_δ-1 = 587 bp, α_2_δ-2 = 425 bp, α_2_δ-3 = 356 bp, α_2_δ-4 = 480 bp. L = ladder.

### 3. α_2_δ-3 expression is widespread in the mouse retina, but strongest in horizontal cells

To determine the cell types expressing α_2_δ-3 in mouse retina, we took advantage of the LacZ insert in knockout and heterozygous mice. After immunostaining against markers of retinal cell types in cryo slices, we performed X-gal staining for cells carrying the LacZ insert. Calbindin - a marker for horizontal cells in mouse retina - showed complete overlap with the X-gal stained cells in the distal inner nuclear layer (INL) (Fig. 3, n = 3 animals). All cells in the distal INL stained for calbindin were also X-gal-positive and vice versa. A very faint X-gal staining was observed in cells in the proximal INL (Fig. 3 A, arrows) and rarely in cells in the ganglion cell layer (Fig. 3 A, arrowhead). We consistently observed this large difference in X-gal staining intensity between horizontal cells and the labeling in proximal INL and ganglion cell layer (GCL). Strong expression of α_2_δ-3 could also be localized to horizontal cells in wholemount immunohistochemical stainings for β-galactosidase and calbindin in developing mouse retina (postnatal day 11, not shown).

**Figure 3:**
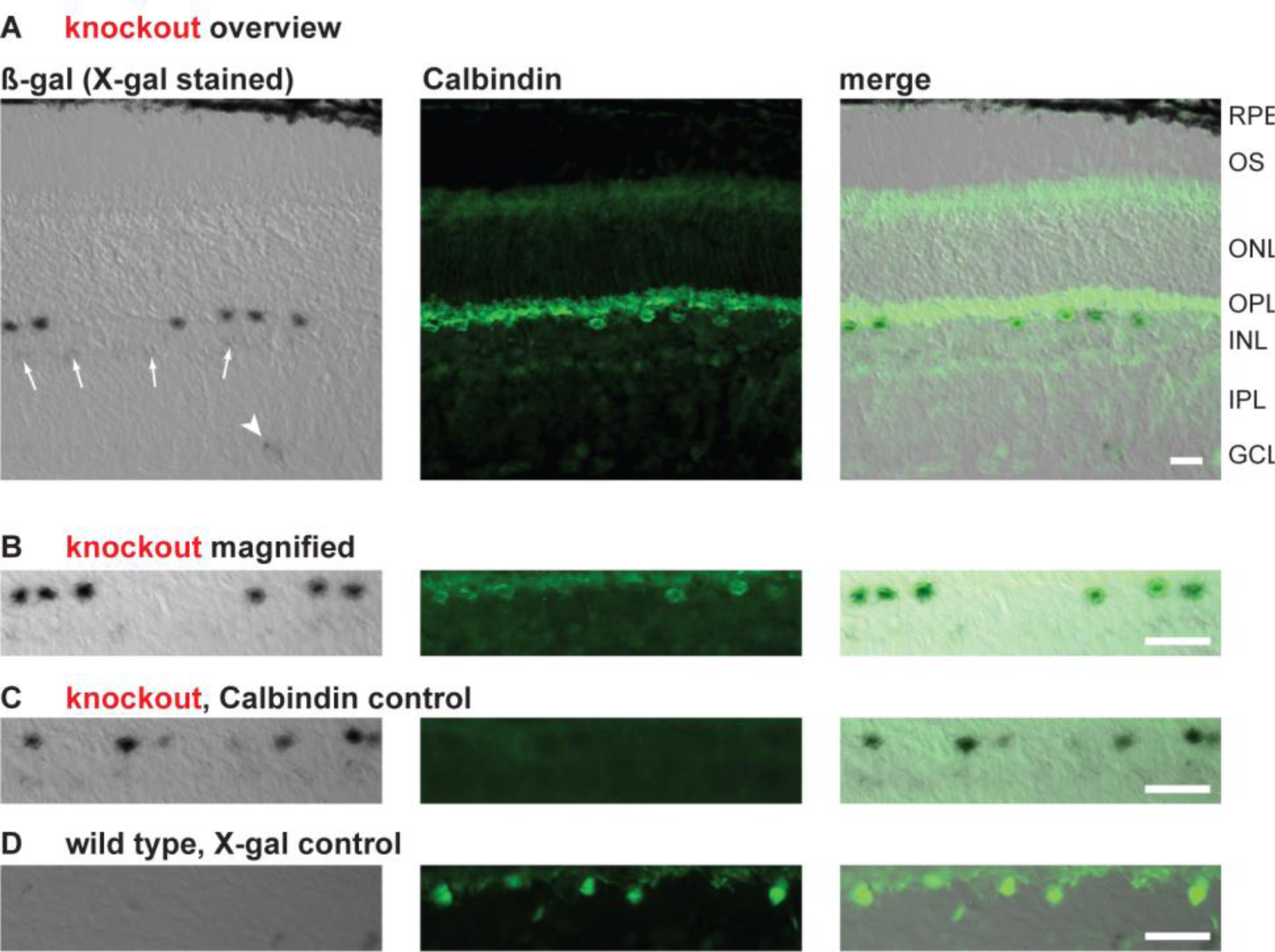
LacZ expression localization in adult mouse retina. Horizontal cells were immunohistochemically stained with antibodies against calbindin in cryoslices of knockout/heterozygous and wild type retina and LacZ reporter expression was revealed with X-gal. **(A)** The strongly LacZ-positive cells showed complete overlap with the calbindin-positive cells in the distal inner nuclear layer (INL). Weak X-gal staining was also observed in cells within the proximal INL (arrows) and some rare cells in the ganglion cell layer (arrowhead). **(B)** Higher magnification of the INL region from panel A. Labeling of cells in the outer INL is very strong. **(C)** The X-gal staining pattern was similar without prior immunohistochemical staining of horizontal cells. **(D)** In wild type controls, no X-gal stained cells could be observed. All scale bars: 20 µm. Abbreviations: RPE = retinal pigment epithelium, OS = photoreceptor outer segments, ONL = outer nuclear layer, OPL = outer plexiform layer, INL = inner nuclear layer, IPL = inner plexiform layer, GCL = ganglion cell layer.

We used immunohistochemical labeling of β-galactosidase (β-gal) in adult mouse retina for easier co-localization of β-gal expression with other cell type markers. Like in X-gal staining, cells in the distal INL were labeled by β-gal-antibody most intensively (Fig.4A), while labeling intensity was weaker in proximal INL and GCL (Fig. 4B+E). We did not find co-localization of the β-gal expression with choline acetyl-transferase (ChAT), a marker for starburst amacrine cells, in the INL (Fig. 4 B-D; n = 3 animals) or in the GCL (not shown). Co-localization of β-gal was found in the majority of Melanopsin-positive ganglion cells in the GCL (Fig. 4 E-G, arrowheads; n = 3 animals) as well as Melanopsin-positive cells in the INL (not shown). Many cells with clear β-gal labeling in the GCL were Melanopsin-negative.

**Figure 4:**
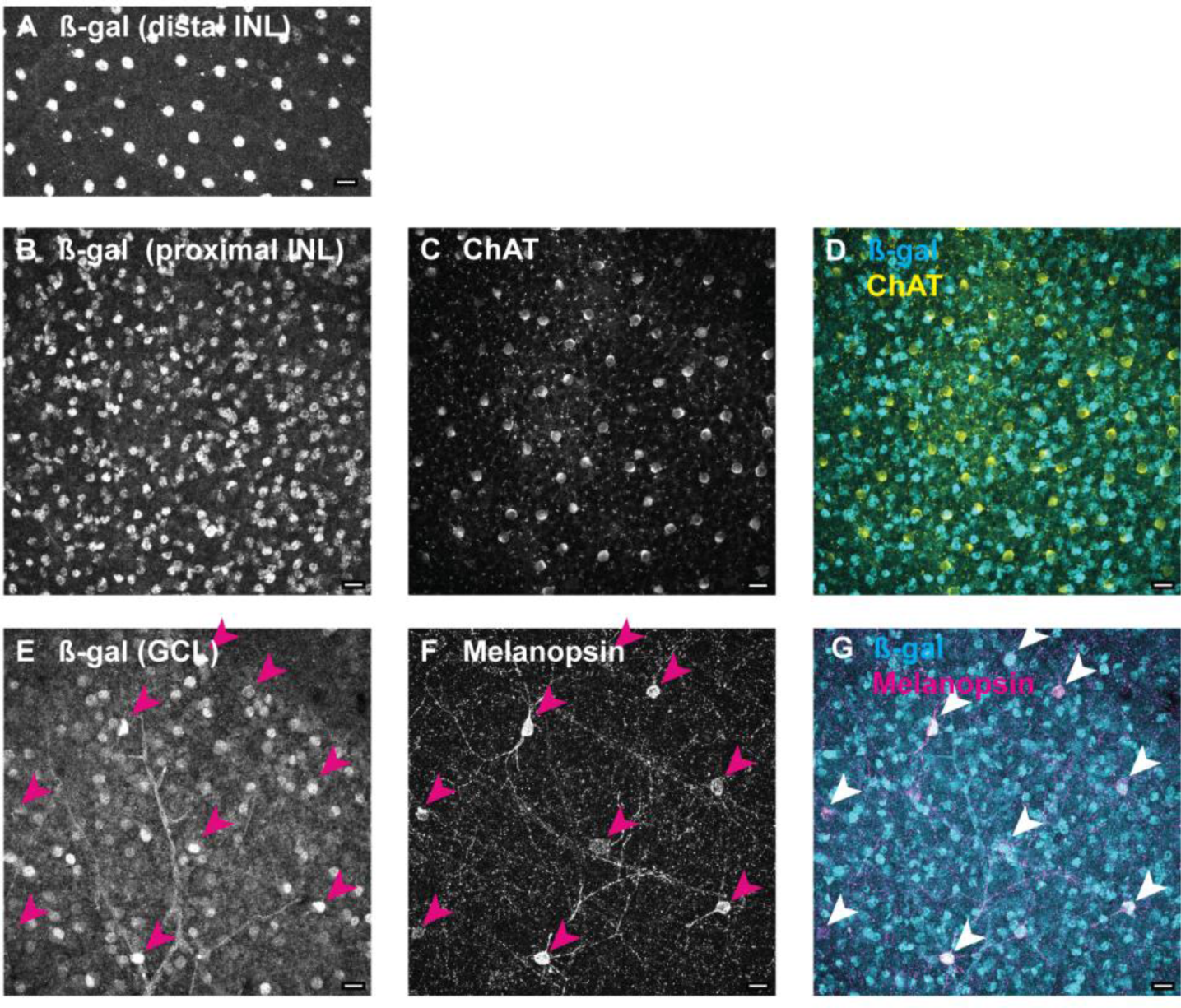
Immunohistochemical localization of β-galactosidase in retinal wholemounts reveals more LacZ-positive cell types. LacZ-positive cells were labeled with an antibody against β-galactosidase and counterstained with markers for different cell types. (**A**) Strong labeling of cells in distal INL was observed, presumably horizontal cells. Staining in proximal INL (**B**) and GCL (**E**) was much weaker (images were acquired with the same settings of the confocal microscope). β-galactosidase staining did not co-localize (white in overlay images) with choline acetyltransferase (ChAT) staining in INL (**B-D**) or GCL (not shown), but with melanopsin staining (**E-G**). Scale bars: 20 µm. Abbreviations: INL = inner nuclear layer, GCL = ganglion cell layer.

### 4. The horizontal cell mosaic is unperturbed by the α_2_δ-3 knockout

To assess possible disturbances of horizontal cell morphology and survival, we labeled horizontal cells (see methods) in wholemount retinas of adult wild type and knockout mice (n = 5 animals each). We covered dorsal and ventral areas between 25% and 75% retinal eccentricity in our analysis (Fig. 5A). Horizontal cell soma positions were determined semi-automatically with a custom Wolfram Mathematica script, and nearest neighbor distances were calculated (25932 cells in wild type and 30093 cells in knockout retinas from n = 5 animals each). Nearest neighbor distance was defined as the distance between the centroid position of a given cell body and the centroid position of the cell body closest to it (Fig. 5B). Nearest neighbor distance was found to increase with retinal eccentricity in both the ventral and the dorsal half of the examined retinas. In the dorsal retina, distances were consistently smaller than in the ventral retina (Fig. 5C). In dorsal retina, average distances were 22.11 ± 2.56 µm (wild type) and 22.03 ± 2.28 µm (knockout). In ventral retina, average distances were 24.29 ± 2.79 µm (wild type) and 24.14 ± 2.57 µm (knockout, means ± standard deviations). There was no statistically significant difference between wild type and knockout in neither dorsal nor ventral retina (p = 0.947, dorsal; p = 0.900, ventral; t-test), while within genotypes the difference between dorsal and ventral was significant (p = 0.0095, wild type; p = 0.033, knockout; paired t-test). The corresponding cell densities in dorsal retina were 1064 ± 209 cells/mm² for wild type and 1081 ± 234 cells/mm² for knockout retinas. Ventral retina densities were 906 ± 190 cells/mm² for wild type and 935 ± 185 cells/mm² for knockout retinas (means ± s.d.). We also looked at the horizontal cell densities as a function of retinal eccentricity but did not find any significant differences between genotypes (not shown). The regularity of the horizontal cell mosaic was assessed by looking at the variability of nearest neighbor distance, which is represented in the distribution of distances (Fig. 5D). A larger variability of nearest neighbor distance would be reflected in a broader distribution, longer tails or larger skewness. We used the standard deviations from the mean distance of each analyzed image as a statistical measure for variability. We found no significant difference in nearest neighbor distance standard deviations between wild type and knockout in neither dorsal nor ventral retina (p = 0.877, dorsal; p = 0.422, ventral; t-test). In summary, we found no effect of the knockout of the α_2_δ-3 gene on horizontal cell number, distribution, and mosaic regularity.

**Figure 5:**
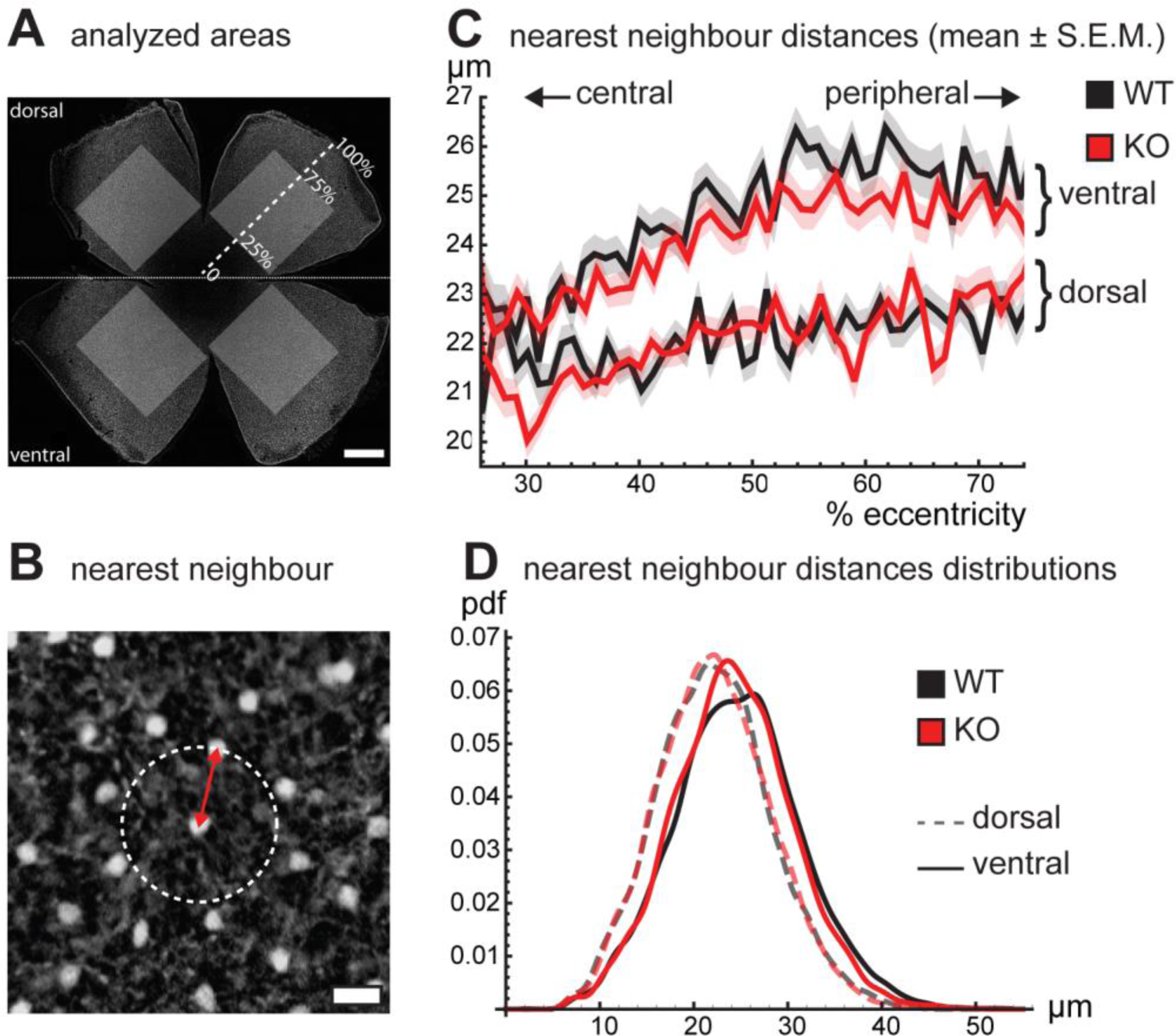
Horizontal cell mosaic analysis. (**A**) Horizontal cells were stained in retinal wholemounts and areas of dorsal and ventral retina from 25% to 75% retinal eccentricity (shaded rectangles; 0% = optic nerve head, 100% = retinal rim) were analyzed. Scale bar: 200 µm. (**B**) Nearest neighbor distances between cells (red arrow = nearest neighbor distance, dotted circle indicates no other cell within same distance; scale bar: 20 µm). (**C**) Nearest neighbor distance plotted as a function of retinal eccentricity shows larger distances between horizontal cells in ventral retina and an overall increase in distances towards the periphery in both wild type and knockout (mean ± SEM). (**D**) Distribution of nearest neighbor distances is very similar between genotypes in either dorsal or ventral retina, indicating a similar regularity of the horizontal cell mosaic. In both genotypes, the distribution in ventral retina is shifted to larger distances (pdf = probability density function). No significant differences of nearest neighbor distances were found between genotypes.

### 5. Currents carried by voltage-gated calcium channels in isolated horizontal cell somata are unchanged

We performed patch-clamp recordings of currents carried by voltage-gated calcium channels (VGCC) from acutely isolated horizontal cell somata in the whole-cell mode to determine an influence of α_2_δ-3 on calcium currents in these cells. Horizontal cell somata were identified by their characteristic morphology (Fig. 6A). Resting membrane potentials of the recorded horizontal cells were −13.06 ± 5.26 mV in wild type and - 12.35 ± 4.14 mV in knockout, and membrane capacitances were 13.38 ± 3.45 pF in wild type and 12.67 ± 2.25 pF in knockout (means ± s.d.). Currents were recorded using 10 mM barium as charge carrier while other conductances through voltage-gated sodium and potassium channels were blocked pharmacologically (see methods).

**Figure 6:**
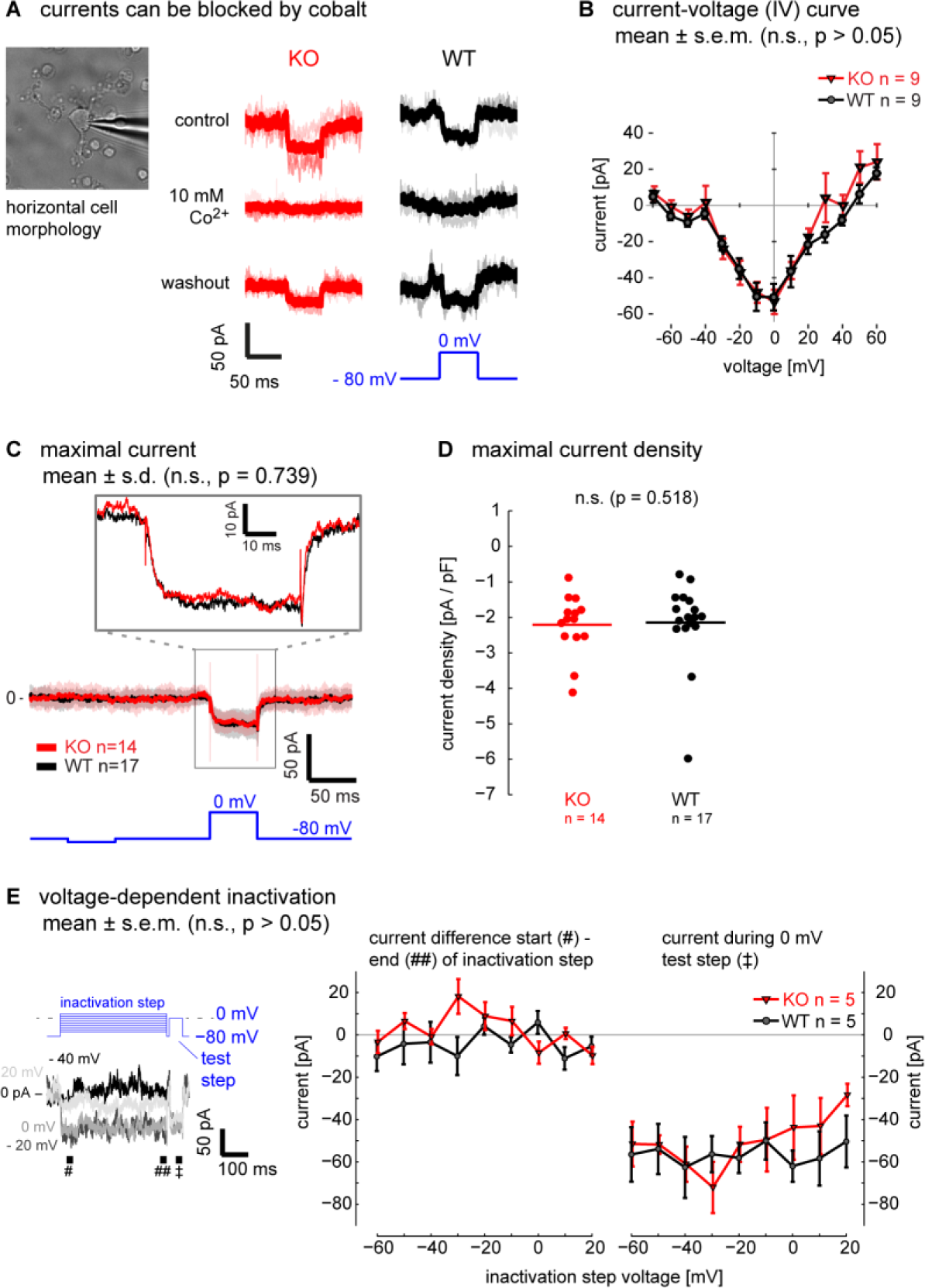
Patch-clamp recordings. (**A**) Acutely isolated horizontal cells were targeted for whole-cell patch-clamp recordings by their characteristic morphology. The currents were confirmed to be carried by voltage-gated calcium channels by blocking with Co^2+^ (thick lines represent averages of 3-4 traces). (**B**) Current-voltage (IV, mean ± s.e.m.) curves of knockout and wild type. The dip at −50 mV could be indicative of T-Type channel activation. There was no significant difference in fitted IV curves half-maximal activation, peak, slope or reversal potential. (**C, D**) The maximal current amplitude to a 0 mV step was the same in knockout and wild type cells, as seen in the averaged traces (**C**, mean ± s.d.) and in the quantification of the maximal current density (**D**, dots = individual cells, horizontal line = mean). (**E**) Both wild type and knockout do not show voltage-dependent inactivation. Inactivation steps from −60 to +20 mV lasted 400 ms followed by a 50 ms test step to 0 mV. Current difference between beginning and end of the inactivation steps and current during the 0 mV step were compared and no significant differences between genotypes were found.

We confirmed that the currents were indeed carried by VGCC by blocking them with a solution where Ba^2+^ was equimolarly replaced by Co^2+^ (Fig. 6A). Current-voltage relationships were tested with 50 ms steps from −70 mV to +60 mV (Fig. 6B). No differences were found between genotypes in peak activation voltage (KO: −9.12 ± 7.69 mV; WT: −8.05 ± 4.99 mV; p = 0.950), half-maximal activation voltage (KO: −21.77 ± 8.19 mV; WT: −20.36 ± 5.97 mV; p = 0.948), slope factor (KO: 8.44 ± 4.03; WT: 9.29 ± 5.78; p = 0.951) or reversal potential (KO: 42.58 ± 7.68; WT: 46.63 ± 7.04; p = 0.222; all parameters derived from fits of IVs; Wilcoxon ranksum tests; n = 9 each). The small negative peak at −50 mV was consistently observed in many cells of both genotypes. Maximal currents were further investigated with a 50 ms step protocol to 0 mV (Fig. 6C). Maximum current amplitudes were −27.29 ± 12.89 pA in wild type (n = 17) and −26.61 ± 8.99 pA in knockout (n = 14, Fig. 6C), the corresponding current densities were −2.14 ± 1.18 pA/pF in wild type and −2.21 ± 0.85 pA/pF in kockout (means ± s.d., Fig. 6D). There was no significant difference between genotypes in maximum current amplitudes (p = 0.739) nor current densities (p = 0.518, Wilcoxon ranksum tests). We looked for voltage-dependent inactivation by applying protocols with 400 ms long depolarizing steps from −80 mV to different target voltages (−60 mV to +20 mV), followed by a 50 ms test step to 0 mV (Fig. 6 E). We observed little to no voltage-dependent inactivation and no differences between genotypes (p > 0.05, Wilcoxon ranksum tests).

### 6. Electroretinography (ERG) shows normal outer retina function

To assess functionality of synaptic transmission in the outer retina, we performed ERG recordings *in vivo* (n = 4 knockout, n = 5 wild type animals).

Under dark-adapted (scotopic) conditions (Fig. 7A), the initial part of the first negative deflection after a light flash (a-wave) is evoked mainly by the rod photoreceptors, whereas the following positive deflection (b-wave) reflects inner retinal activity, including depolarizing (ON) bipolar cells. Note that under scotopic conditions, from 10^-2^ cd*s/m² onward (arrow in Fig. 7A), cones also get activated. Under light-adapted (photopic) conditions (Fig. 7B) the b-wave relies on activity driven by the cone system, with the rod system being saturated by the background illumination. Scotopic single-flash ERG recorded from α_2_δ-3 knockout mice did not show significantly different b-wave amplitudes or latencies from those in wild type animals (Fig. 7C). There were also no differences in the b-wave amplitudes or latencies of photopic single-flash ERG responses of α_2_δ-3 knockout and wild type mice (Fig. 7D).

**Figure 7:**
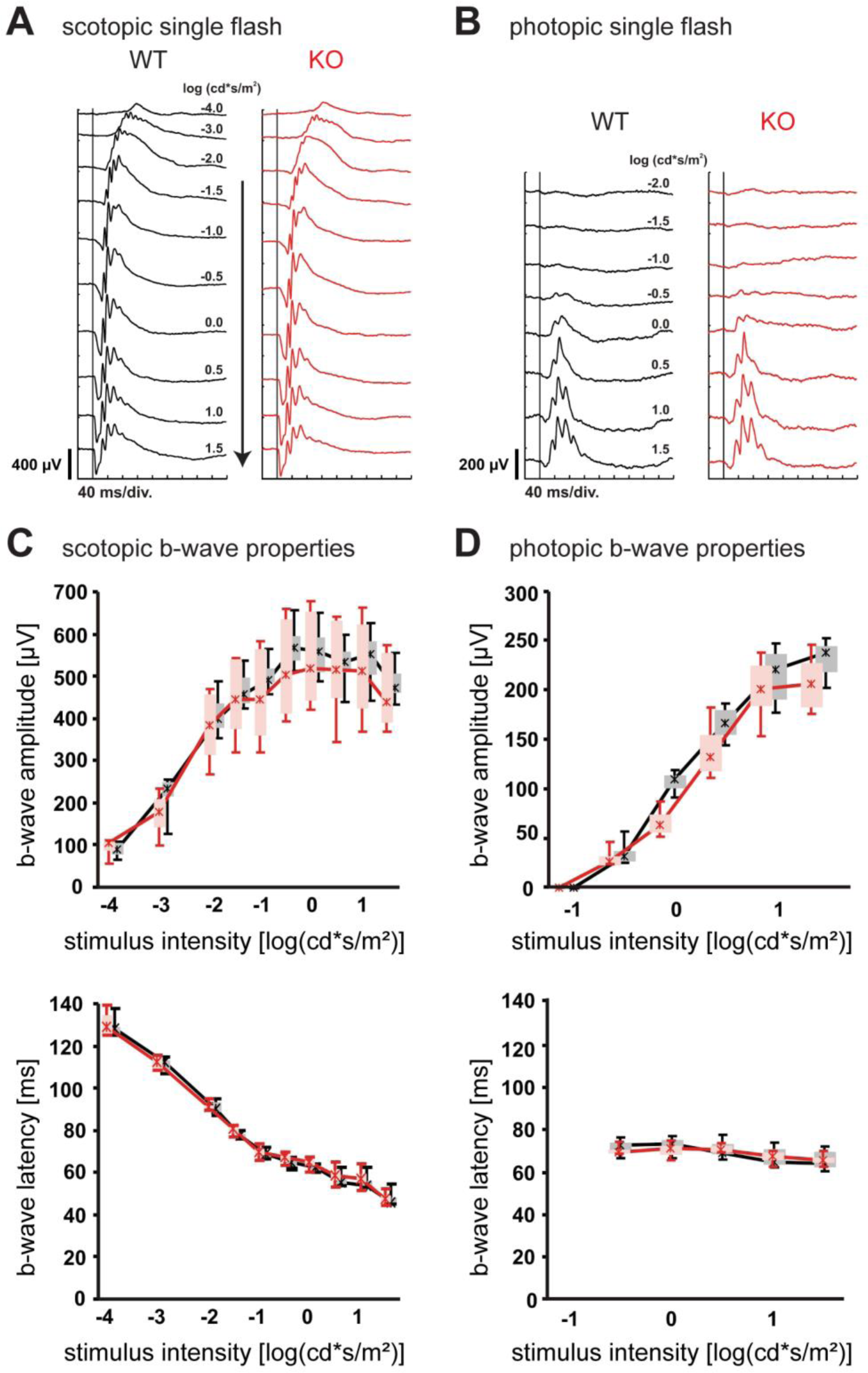
ERG *in vivo* functional analysis. (**A**) Representative single-flash ERG intensity series of wild type (black) and knockout mice (red) under scotopic conditions (arrow indicates intensities with cone activation) and (**B**) under photopic conditions. (**C**) Box-and-whisker plots showing single-flash b-wave amplitudes and b-wave latencies under scotopic conditions and (**D**) under photopic conditions of wild type and knockout mice. Asterisks = medians, connected by solid lines; boxes = 25% to 75% quantile range; whiskers = 5% and 95% quantiles.

### 7. Multi-electrode array (MEA) recordings from retinal ganglion cells reveal several subtle phenotypes of the α_2_δ-3 knockout retina

In order to investigate perturbances of retinal processing and possible effects downstream of bipolar cells, we isolated retinas from wild type and knockout mice and recorded the activity of retinal ganglion cells with micro-electrode arrays (MEAs). During the recording, light stimuli were presented that covered scotopic, mesopic, and photopic intensity levels (see methods). The same set of stimuli was presented twice at each ambient luminance level. A legend of the presented stimuli is shown at the bottom of Fig. 8A.

**Figure 8:**
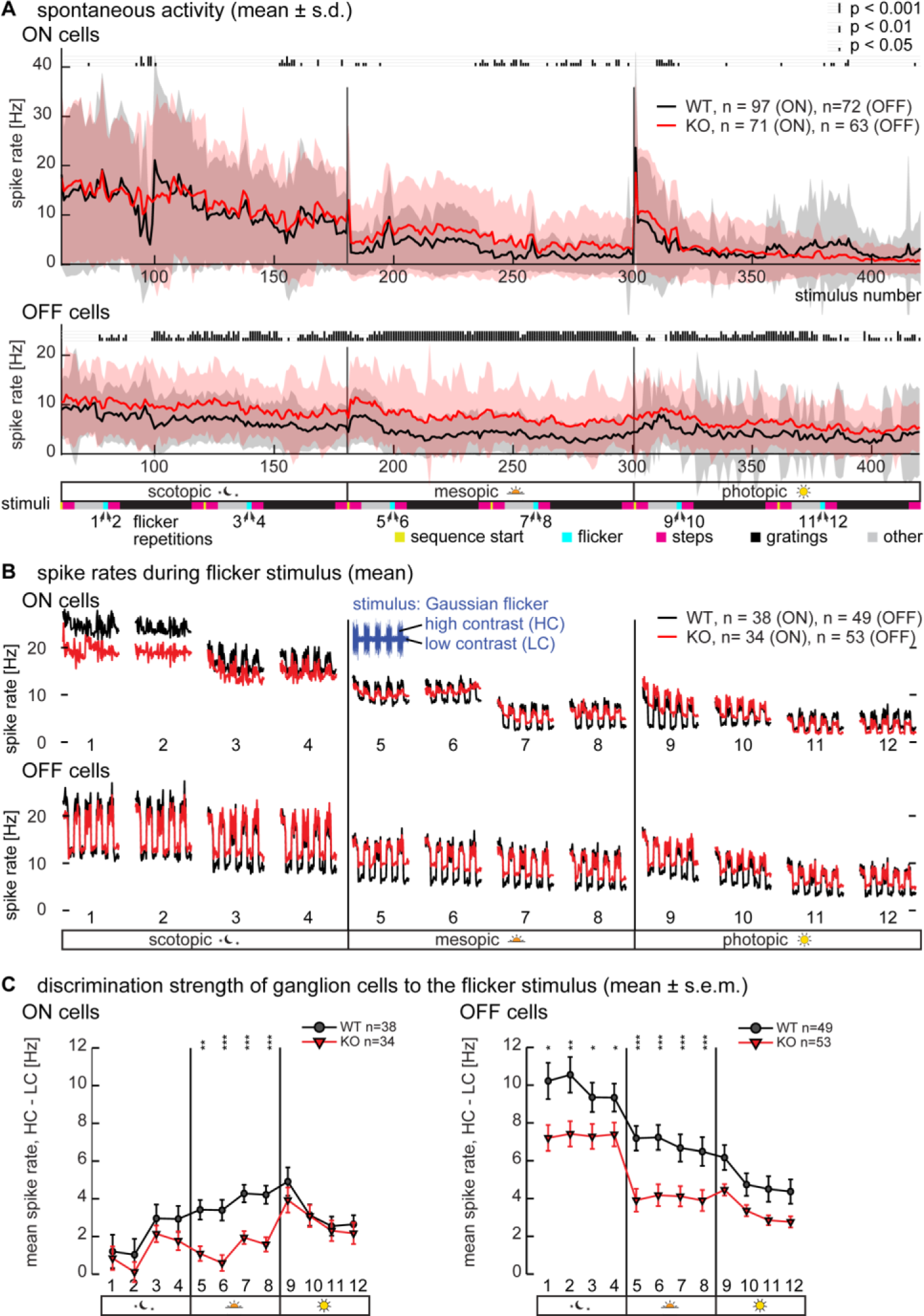
Multi-electrode array recordings of ganglion cell spontaneous activity and responses to Gaussian white noise flicker. Ganglion cell activity was recorded with multi-electrode arrays while the retina was exposed to a sequence of visual stimuli, repeated at each ambient luminance level. Each visual stimulus was preceded by 1 second of uniform background gray (‘128’). This time window was used to calculate spontaneous activity. (**A**) Spontaneous activity in ON and OFF ganglion cells (polarities defined by STA) over the whole experiment (stimulus durations were 12 seconds to 3 minutes; stimuli 1 to 60 were considered as “settling down period” and not analyzed). ON cell spontaneous activity (upper panel) differed mainly during mesopic conditions while OFF cells (lower panel) in knockout exhibited an increased spontaneous spike rate over all luminance levels (height of the vertical bars on top of the panels indicates p-values). Vertical lines indicate switches to the next ambient luminance level. Stimuli are indicated below: Yellow – sequence start, magenta – full field steps, cyan – Gaussian white noise flicker, black – drifting gratings, gray – other. (**B**) Mean spike rates during the full-field Gaussian white noise flicker stimulus with interleaved high-contrast (HC) and low-contrast (LC) episodes. For Gaussian flicker, only cells with a clear STA to LC episodes of at least one stimulus repetition per luminance level were analysed. In general, spike rates during HC episodes were similar between genotypes, while spike rates during LC episodes were higher in knockout ON and OFF ganglion cells. (**C**) The difference in spike rate between HC and LC episodes was calculated as a measure of discrimination strength of the contrast modulation. ON cells (left panel) showed a clear difference in discrimination strength only during mesopic conditions, while OFF cells (right panel) had significantly different discrimination strength during mesopic and scotopic conditions.

#### Spontaneous activity

Spontaneous activity was determined as the spike rates during the first second of each stimulus, during which a uniform gray (‘128’) background was always presented. Note, that due to the short times between stimuli in our experiments (around 3 seconds), the spontaneous activity preceding a stimulus might still be influenced by the previous stimulus. Spontaneous spiking was highest during scotopic ambient luminance and decreased over the time course of the experiment in ON cells (Fig. 8 A, top) and less so in OFF cells (Fig. 8 A, bottom). ON cells had different spontaneous spike rates in wild type and knockout retina mostly during mesopic ambient luminance (Fig. 8 A, upper panel, statistical significance indicated on top with bars of different heights, Wilcoxon ranksum tests; KO: n = 71, WT: n = 97 cells). Interestingly, OFF ganglion cells in knockout retina had a markedly higher spontaneous spike rate than in wild type retina over most of the time course of the experiment (Fig. 8 A, lower panel; KO: n = 63, WT: n = 72 cells).

#### Gaussian white noise stimulus

One of our stimuli was a full-field Gaussian white noise (“flicker”) stimulus. Each flicker stimulus consisted of episodes of high-contrast flicker interleaved with episodes of low-contrast flicker (Fig. 8B). The stimulus was repeated four times at each ambient luminance level, for a total of 12 repetitions (marked 1 through 12 in the legend at the bottom of Fig. 8A). Spike rates of ganglion cells increased during high-contrast episodes, reflecting the stronger drive by the higher stimulus contrast. During low-contrast episodes, spike rates decreased and dropped to approximately the spontaneous rate before stimulus onset. In both ON and OFF cells, the spike rates during high-contrast seemed more similar between wild type and knockout, than spike rates during low-contrast.

For the analysis, we subtracted the spike rates during low-contrast episodes from the high-contrast spike rates (Fig. 8C) for each stimulus repetition. We refer to this difference as the discrimination strength of a ganglion cell to the flicker stimulus. We only included cells in the analysis that had a clear spike-triggered average (STA) derived from the low-contrast episodes of at least one stimulus repetition per light level to ensure that the spikes are stimulus-driven (ON cells: KO n = 34; WT n = 38. OFF cells: KO n = 53; WT n = 49). In ON ganglion cells (Fig. 8C left), we did not observe significant differences between wild type and knockout mice in discrimination strength during scotopic and photopic conditions (0.413 < p < 0.992). However, in mesopic conditions the discrimination strength was significantly lower in knockout compared to wild type (p < 0.01 for first, p < 0.001 for other three repetitions). In OFF ganglion cells (Fig. 8C right), we observed significantly reduced discrimination strength in knockout during scotopic (p = 0.012, p = 0.005, p = 0.037, p = 0.039) and mesopic conditions (p < 0.001, all repetitions), but not in photopic conditions (0.053 < p < 0.487; Wilcoxon ranksum tests).

#### Drifting sinusoidal grating stimulus

Looking carefully at the spontaneous activity plots, we observed peculiar jumps in spontaneous activity in ON ganglion cells during drifting sinusoidal grating stimuli in scotopic luminance (Fig. 8A, top, before stimulus number 100 and after stimulus 150). To investigate this further, we grouped ganglion cells based on their spontaneous activity following sinusoidal grating stimuli in scotopic luminance (Fig. 9A). We clustered the ganglion cells by a k-means algorithm (Matlab) to avoid any subjective selection bias. Most clusters were approximately equally populated by wild type and knockout cells. However, we found two clusters that were populated mostly by either wild type cells (cluster 4) or by knockout cells (cluster 8), and both exhibited a clear jump in spontaneous activity in scotopic luminance. These two clusters robustly emerged when running the k-means algorithm with a value of k (= number of clusters) between 7 and 12. Fig. 9A shows the distribution for k = 8 clusters. Four clusters consisted exclusively of ON cells (1, 4, 5, 8), while the other four clusters were mixed. While both clusters 4 and 8 exhibited a jump in spontaneous activity, this jump happened after presentation of gratings of different spatial properties. This is evident in the spontaneous activities of the individual cells of clusters 4 and 8 during scotopic luminance shown in Fig. 9B. Spike rates decreased after the presentation of gratings with low spatial periods, sometimes exhibiting a marked drop. Common to all cells in clusters 4 and 8 was a sharp increase in spontaneous spike rate: in cluster 8, this increase happened after presentation of the first 2000 µm grating, while cluster 4 shows a similar jump in spontaneous activity in between the 500 µm and 1000 µm gratings. In both clusters the increased spontaneous activity was maintained between presentations of the gratings with large spatial periods and only occurred in scotopic luminance. A natural question is, whether the change in spontaneous activity after the presentation of certain gratings simply reflects a persisting elevated activity of the cells during those gratings, i.e. if our observation can be explained by a lingering effect of responses to the stimulus itself. Fig. 9C shows example responses after aligning (phase shifting) these responses by cross-correlation with the sinusoidal grating stimulus (phase-shifts in responses of different ganglion cells stem from different receptive field positions relative to the grating stimulus). Of cluster 4, only wild type cells were considered for further analysis. We averaged spike rates of cells within each cluster and determined the following response parameters with or without subtracting the spontaneous spike rate (baseline): minimal spike rate, maximal spike rate, mean spike rate, median spike rate and response amplitude (max-min). We compared the parameters of gratings of the same temporal frequency, but with different spatial periods. Fig. 9D plots significant changes in a heatmap (Wilcoxon ranksum test). For example, the top left entry indicates the level of significance when comparing the presentations of 0.25 Hz gratings with 100 versus 200 µm spatial periods. Of all parameters examined, only the baseline-subtracted mean spike rate yielded significant differences between gratings of different spatial properties in the range with a change in the spontaneous spike rate (Fig. 9D, left; spatial period range marked with yellow rectangle). In all other parameters, only the strong changes between 200 µm and 500 µm gratings yielded systematically significant differences, shown here for the parameter “amplitude” of clusters 4 and 8 (Fig. 9D, right). The situation was similar for all eight clusters (not shown). In summary, the change of spontaneous activity after the presentation of certain gratings is hardly reflected by changed response properties to the gratings themselves.

**Figure 9:**
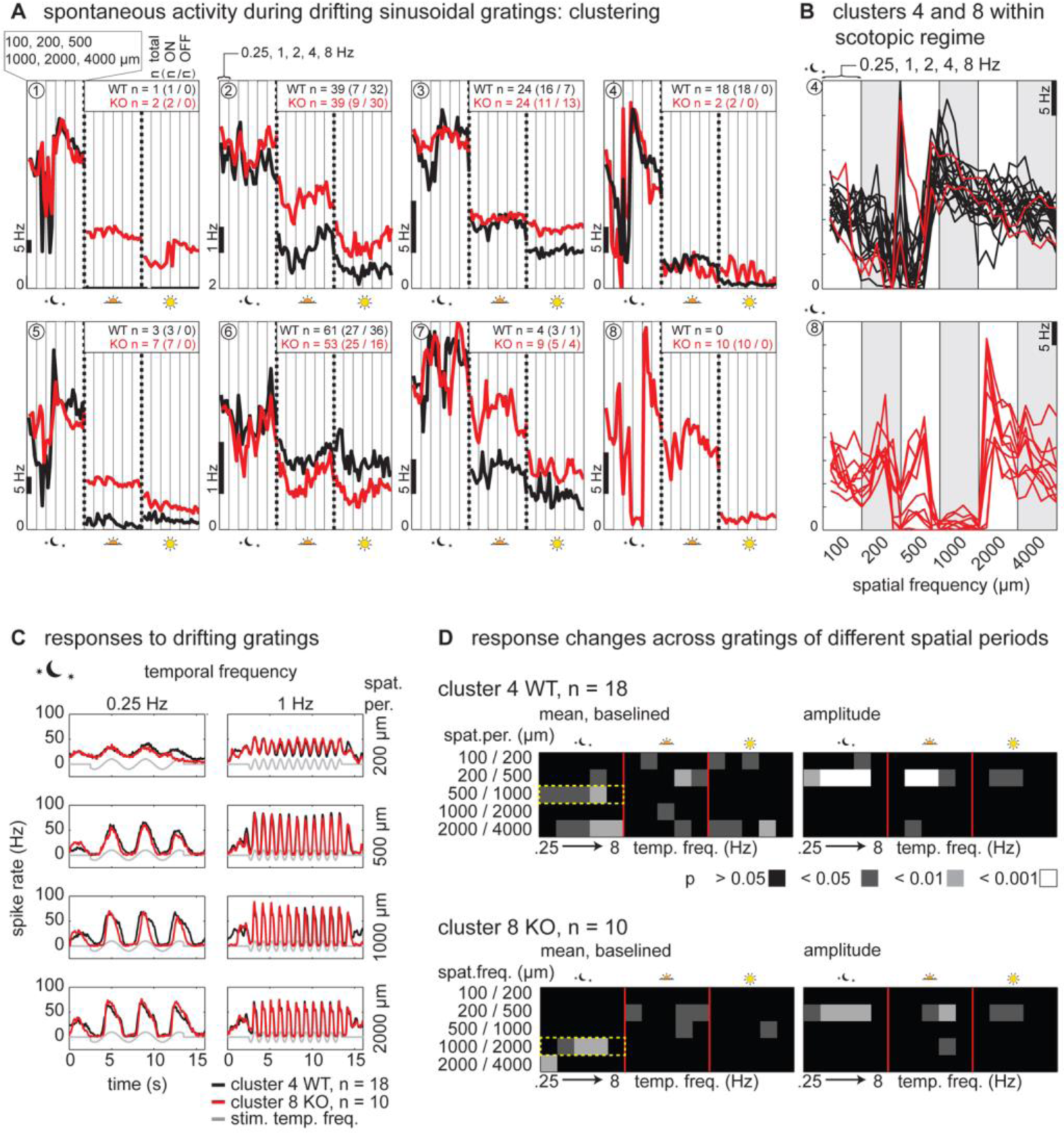
Multi-electrode array recordings of ganglion cell responses to drifting sinusoidal gratings. Ganglion cells were k-means-clustered based on their spontaneous activity following sinusoidal grating stimuli in scotopic luminance. (**A**) Mean spontaneous activity of eight ganglion cell clusters. Four clusters were exclusively ON cells (1, 4, 5, 8), while the other four clusters were mixed (n numbers: total (ON cells/OFF cells)). Most clusters contained wild type and knockout ganglion cells, only clusters 4 and 8 were more specific. Columns mark spatial periods, each containing all temporal frequencies (see labels above clusters 1 and 2). Vertical dotted lines indicate the next luminance level. (**B**) Spontaneous activity of individual cells of clusters 4 (top) and 8 (bottom) during scotopic luminance. Spike rates decreased during the low spatial periods, with a sharp increase during 500 µm gratings (cluster 4) or after the presentation of the 2000 µm gratings (cluster 8). (**C**) Averaged response of cluster 4 (WT only) and cluster 8 to a subset of grating stimuli. There is a marked increase in spike rate modulation (amplitude) in the responses to the 500 µm gratings compared to the 200 µm gratings. (**D**) Only the baselined mean spike rate (left panels) differed significantly at the junctions of grating spatial periods where we saw spontaneous activity jumps (yellow rectangles). Amplitudes (right panels) showed significant differences mainly at the junction between 200 µm and 500 µm gratings.

#### Full-field step stimulus

We next analyzed the responses to a full-field contrast step stimulus. The averaged responses were very similar in OFF ganglion cells from wild type and knockout at all three luminance levels (Fig. 10A). Only subtle differences between genotypes in the positive contrast steps (going ‘brighter’) at scotopic and mesopic luminance could be observed. In OFF ganglion cells of the knockout, response peaks were partly shifted in time (arrows) and response amplitudes were increased (arrowheads). Note that classical ON-OFF cells would almost unequivocally fall into the OFF category in our classification by their STA polarity (Tikidji-Hamburyan et al., 2014).

**Figure 10:**
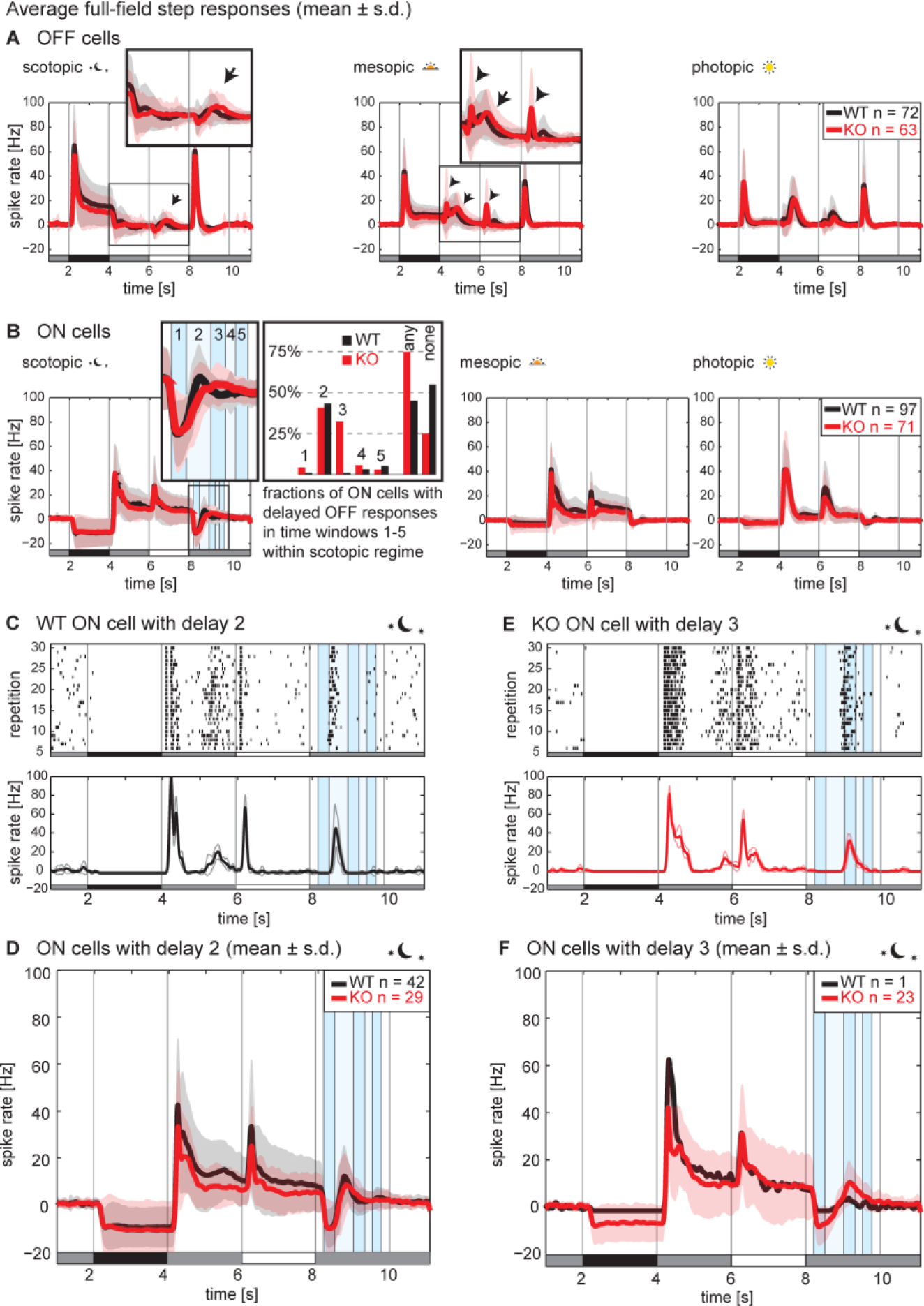
Multi-electrode array recordings of ganglion cell responses to full-field contrast steps. Ganglion cells were stimulated with full-field steps: gray → black → gray → white → gray (see bottom of each panel). (**A**) Averaged responses (mean ± s.d.) of OFF cells from wild type and knockout at all three luminance levels. Response peaks to anti-preferred contrast steps were partly delayed (arrows) and response amplitudes were increased (arrowheads) in scotopic and mesopic luminance. (**B**) Responses of ON cells of wild type and knockout were very similar in mesopic and photopic luminance levels. In scotopic luminance, responses to the last step were delayed. We observed several subpopulations characterized by the delay timings (see standard deviations in inset). Subpopulations were similar in wild type and knockout, except for cells with a delayed response in time window 3, which was almost exclusively found in knockout (bar graph). (**C**) Example scotopic response of a WT ON cell with delay 2 (top: raster plot of all 30 repetitions; bottom: averaged spike rate; thick line = mean, thin lines = s.d.). (**D**) Averaged scotopic response of all cells with a delayed response in time window 2 (mean ± s.d.). (**E**) Example scotopic response of a KO ON cell with delay 3 (top: raster plot of all 30 repetitions; bottom: averaged spike rate; thick line = mean, thin lines = s.d.). (**F**) Averaged scotopic response of all cells with a delayed response in time window 3 (mean ± s.d.).

ON cells of wild type and knockout also responded very similarly in mesopic and photopic luminance levels (Fig. 10B). However, in scotopic luminance, upon return to the grey background after the white step, we observed different delays in response peaks. The averaged responses revealed that there are several subpopulations with different properties, as it appears from the peaks of the standard deviations of the responses (inset). To investigate this further, subpopulations of cells were defined by manually classifying response peaks in five time windows after the offset of the white flash: 0.2 – 0.5 s, 0.5 – 1.0 s, 1.0 – 1.3 s, 1.3 – 1.5 s and 1.5 – 1.75 s. This yielded a differential distribution pattern in wild type and knockout (bar graph in Fig. 10B). 75% of ON cells in knockout had such responses versus 45% of the wild type cells. Strikingly, almost all wild type cells with a response fell into subpopulation ‘2’ (response peak between 0.5 and 1 s). Subpopulation ‘3’ (1 to 1.3 s) was almost exclusively found in knockout retinas. Fig. 10C and E illustrates the marked timing difference in these responses with two representative cells belonging to subpopulations ‘2’ and ‘3’ (spike raster on top, spike rate at the bottom). Fig. 10D and F show the population averages of these two populations.

## Discussion

We found especially strong expression of the voltage-gated calcium channel (VGCC) subunit α_2_δ-3 in horizontal cells in the mouse retina. Yet, a knockout of α_2_δ-3 did not lead to changes of the horizontal cell mosaic or of voltage-gated calcium channel currents within horizontal cell somata. Outer retinal function measured by electroretinograms was normal, as was the optokinetic reflex behavior in α_2_δ-3 knockout animals. In ganglion cells however, we could see a number of changes in response properties to different kinds of visual stimuli, which were mostly restricted to scotopic or mesopic ambient luminance levels of our stimulation paradigm.

### α_2_δ mRNA expression

Gene or protein expression of α_2_δ-1 (Eroglu et al., 2009; Huang et al., 2013), α_2_δ-3 (Nakajima et al., 2009; Pérez de Sevilla Müller et al., 2015) and α_2_δ-4 (Wycisk et al., 2006; Mercer et al., 2011; de Sevilla Muller et al., 2013; Thoreson et al., 2013) have been reported in vertebrate retina. To the best of our knowledge, expression of α_2_δ-2 in adult retina, or expression of any of the four α_2_δ subunits during mouse retinal development have not been demonstrated. We found expression of all α_2_δ genes in adult as well as in developing mouse retina, from at least postnatal day 3 onwards (Fig. 2). This raises the possibility for a function of the α_2_δ subunits in synaptogenesis (Kurshan et al., 2009) in mouse retina during development.

### α_2_δ-3 localization

It has previously been reported that α_2_δ-3 is expressed in ON bipolar cells (Nakajima et al., 2009), photoreceptors, bipolar, amacrine and ganglion cells but not horizontal cells (Pérez de Sevilla Müller et al., 2015). Yet, we found very prominent LacZ staining in horizontal cells, driven by the endogenous α_2_δ-3 promotor (Fig. 3). The extraordinary strength of the horizontal cell labeling by the LacZ reporter we observed is completely in line with the α_2_δ-3 gene expression values found in the microarray data of Siegert et al. (2012) (the expression profile can be found online by selecting “*Cacna2d3”* at http://www.fmi.ch/roska.data/index.php). Therefore we believe that the clear antibody labeling of α_2_δ-3 protein in the outer plexiform layer that has been reported by Pérez de Sevilla Müller et al. (2015) stems at least in large part from horizontal cells. The absence of this labeling from cone pedicles would thus suggest localization of α_2_δ-3 only in horizontal cell processes contacting rods. The data of Siegert et al. (2012) also supports our finding of LacZ reporter labelling in melanopsin ganglion cells (Fig. 4). Interestingly, their microarray data shows expression of α_2_δ-3 also in A-II and maybe other gylcinergic amacrine cells, which was also reported by Pérez de Sevilla Müller et al. (2015). This would be consistent with our observation of LacZ labeling in the proximal INL (Fig. 3+4).

### Retina morphology and the horizontal cell mosaic

The mouse retina only contains one type of horizontal cell of the axon-bearing b-type (Peichl and Gonzalez-Soriano, 1994) which form a regular mosaic (Wassle and Riemann, 1978). Mosaic formation is at least in part controlled by repulsive homotypic interactions between neighboring horizontal cells (Poche et al., 2008; Huckfeldt et al., 2009). It has been shown that α_2_δ subunits have a function in synaptogenesis and synaptic stabilization (Eroglu et al., 2009). The unperturbed regularity of the mosaic and spacing of horizontal cells (Fig. 5) can be interpreted as an indication of normal outer retinal development and suggests no impact on horizontal cell survival by the α_2_δ-3 knockout. However, the connectivity and ultra-structure of the triad synapses should be investigated more closely to determine potential effects on synaptogenesis (Reese et al., 2005) and horizontal cell connectivity in detail.

### Voltage-gated calcium channel currents in horizontal cell somata

The α_2_δ subunits are thought to enhance trafficking of VGCC to the membrane. Yet we could not detect a difference in maximum current amplitude or density in horizontal cell somata of α_2_δ-3 knockout nor in the parameters of the current-voltage relationships or the voltage-dependent inactivation (Fig. 6). It is unknown whether α_2_δ-3 interacts with all VGCC α_1_ subunits of horizontal cells, namely L-, P/Q- and N-Type (Schubert et al., 2006; Liu et al., 2013). It is also not known if there is more than one α_2_δ isoform expressed in horizontal cells. Differential regulation of either α_1_ or α_2_δ subunits could compensate for the knockout of α_2_δ-3, rescuing horizontal cells from having a pronounced change in voltage-gated calcium channel properties. The other intriguing possibility is the putative localization of α_2_δ-3 on the rod-contacting axonal processes. Any changes on calcium currents in the axonal compartment would likely not show up in our somatic recordings.

### Outer retina function and visual reflex behavior

Horizontal cells form reciprocal synapses with photoreceptors and this feedback is thought to influence spatial (Thoreson and Mangel, 2012; Szikra et al., 2014) as well as temporal processing properties of the retina (Pandarinath et al., 2010). If the α_2_δ-3 knockout had an impact on feedback, it could affect the strength or kinetics of synaptic transmission from photoreceptors also to bipolar cells, and be reflected in the properties of bipolar cell responses. Our electroretinographic (ERG) recordings did not show an effect of the α_2_δ-3 knockout on latencies or amplitudes of the b-wave (indicative of depolarizing bipolar cell activation), in neither scotopic nor photopic conditions (Fig. 7). These results indicate that α_2_δ-3 subunits are not essential for photoreceptor responses or the synaptic signal transmission to depolarizing bipolar cells.

As a measure for overall retinal functionality we tested the α_2_δ-3 knockout animals for their optokinetic reflex behavior. Tracking behavior in our virtual optokinetic drum experiments was found in most knockout animals (Fig. 1), underlining general retinal functionality and suggesting meaningful output of α_2_δ-3 knockout retina in this simple visual reflex paradigm.

### Retinal processing

Our micro-electrode array (MEA) recordings of retinal ganglion cells revealed several subtle differences between wild type and α_2_δ-3 knockout retina. Some of the effects appear rather general, such as the average spontaneous spike rate which was elevated throughout the experiment (covering all luminance levels) in OFF ganglion cells, but not in ON cells (Fig. 8A). The compression of discrimination strength between high- and low-contrast flicker stimuli (Fig. 8B+C) was restricted to scotopic (ON cells) or scotopic and mesopic luminance levels (OFF cells). This compression seemed to be caused mainly by an elevation of spike rates to the low contrast-condition, while the spike rates during high contrast were largely similar. The restriction of effects to lower luminance levels was a feature we observed consistently.

While OFF cells showed only minor differences in our other stimuli (Fig. 10A), there were subsets of ON cells with changed response profiles which we found only in α_2_δ-3 knockout retinas. We saw a jump in spontaneous spike rates during drifting grating stimuli in both wild type and knockout ON cells, reflecting changed activity after grating stimuli of certain spatial properties. However, these jumps occurred after gratings of different spatial properties in wild type and knockout ganglion cells. It is tempting to speculate that the two clusters of cells with an abrupt change in spontaneous spike rates (Fig. 9B) reflect a change in spatial processing of the involved circuitry. We do not know, however, what brought about this change in spontaneous spike rates, as responses to the preceding grating stimuli themselves did not show any abrupt changes in both clusters. The only difference we found (a change in baseline-adjusted mean spike rates, Fig. 9D) is likely due to the shift in the spontaneous spike rate itself, as this determines the baseline value and did not show in the non-baselined data (not shown). We didn’t observe any systematic differences depending on the temporal properties of the grating stimuli we used.

The responses within different time windows of our full-field step stimuli (Fig. 10B), on the other hand, can be seen as a change of temporal properties. We do not know the physiological significance of these responses, as they only appear on the return to the background mean luminance and most have a very long temporal scale (> 500 ms). We refer to this type of responses as delayed responses. In general, we found delayed responses to be much more abundant in ganglion cells of knockout retina. This could indicate an emergence of this kind of responses in ganglion cells, rather than a shift in the delay time in the same ganglion cell types. It could also indicate a different distribution of the abundance of single ganglion cell types in the knockout, as we cannot rule out sampling artifacts of the MEA technique, i.e. we likely record from different numbers of single ganglion cell types in each recording.

What could cause the differences in retinal ganglion cell responses which we described? Due to the widespread expression of α_2_δ-3 in various cell types, interpretation of the found phenotypes has to be undertaken with care. As a subunit of voltage-gated calcium channels, the α_2_δ-3 knockout is most likely to cause a loss-of-function kind of phenotype, affecting synaptic release in the respective cells. The increased spontaneous spike rate in OFF ganglion cells might be caused by reduced inhibitory transmission somewhere in the circuitry. It is unlikely that the compression of discrimination strength in our Gaussian white noise stimulus was caused by the α_2_δ-3 knockout in horizontal cells, since horizontal cells do not play a role in contrast gain control (Beaudoin et al., 2007) or contrast adaptation (Baccus and Meister, 2002). The shift in spontaneous spike rate during drifting sinusoidal gratings of certain spatial properties could be indicative of a change in receptive field size in one or several cell types. Finally, the abundance of delayed responses and their delay timings in our full-field step stimuli might be caused by a prolonged inhibitory input or by a change in the interaction between the ON and OFF pathways. It is also possible that some of the changes are caused by ganglion cell-intrinsic mechanisms, since α_2_δ-3 expression in the ganglion cell layer seemed low but was widespread. Interestingly, the majority of changes we observed were restricted to brightness regimes with rod activity (scotopic and mesopic), raising the possibility of a role of α_2_δ-3 in retinal circuitries carrying rod signals.

### Horizontal cell function of α_2_δ-3 and other possible mechanisms

It remains open how α_2_δ-3 acts within VGCC complexes in horizontal cells and if it contributes to feedback from horizontal cells to photoreceptors (Thoreson and Mangel, 2012). Horizontal cells have two functionally separated cellular compartments that were thought to be electrically isolated from each other (Nelson et al., 1975): somato-dendritic processes contact cone pedicles, axonal arborizations contact rod spherules (Kolb, 1970, 1974). Signals can, however, spread from the dendritic compartment down this axon and impact the rod circuitry (Trumpler et al., 2008), creating a form of surround inhibition by sign-inverting activation of rod synapses (Szikra et al., 2014). Localization of α_2_δ-3 within the axonal compartment of horizontal cells could at least in part explain the restriction of the phenotypes we found in our MEA recordings to scotopic and mesopic regimes. Horizontal cells might thus also influence the balance between rod and cone activity by utilizing different molecular machinery for feedback in each compartment, involving α_2_δ-3 (rods) or not involving α_2_δ-3 (cones).

Considering a putative disturbance of the rod circuitry, A-II amacrine cells would also be a good candidate for some of the changes we observed in non-photopic regimes. Indeed, glycinergic amacrine cells were reported to express α_2_δ-3 (Siegert et al., 2012; Pérez de Sevilla Müller et al., 2015).

A cell type-specific knockout of α_2_δ-3 in horizontal cells or A-II amacrine cells would help to elucidate the specific effects of this isoform in certain aspects of retinal processing. Especially the possible role of α_2_δ-3 in horizontal cell feedback to rods versus feedback to cones could be interesting to investigate.

## Acknowledgements

The authors thank Sebastian Ströh and Karin Dedek for the introduction to isolated horizontal cell patch-clamp techniques, Nadine Ortner for support on fitting IV curves and Jutta Engel and Timm Schubert for fruitful discussions.

## Role of authors

Designed study: HS, AP, MS, TAM. Performed Experiments: HS, VS, BB, MGG. Analyzed data: HS, VS, BB, ATH, MGG. Wrote paper: HS, VS, BB, TAM.

## Resources cited

B6.129P2-*Cacna2d3^tm1Dgen^*/J mice (The Jackson Laboratory, RRID: IMSR_JAX:005780)

mouse monoclonal anti-calbindin antibody (Swant Cat# 300 RRID:AB_10000347)

rabbit polyclonal anti-calbindin antibody (Swant Cat# CB 38 RRID:AB_10000340)

chicken polyclonal anti-neurofilament H antibody (Abcam Cat# ab4680, RRID:AB_304560)

goat polyclonal anti-choline acetyltransferase antibody (Millipore Cat# AB144P, RRID:AB_2079751)

rabbit polyclonal anti-melanopsin antibody (Advanced Targeting Systems Cat# AB-N39, RRID:AB_1608076)

chicken polyclonal anti-β-galactosidase antibody (Abcam Cat# ab9361, RRID:AB_307210)

Fiji (RRID:SciRes_000137) PatchMaster (v2×69, RRID:SciRes_000168, HEKA)

